# The Landscape of Human STR Variation

**DOI:** 10.1101/004671

**Authors:** Thomas Willems, Melissa Gymrek, Gareth Highnam, The 1000 Genomes Project, David Mittelman, Yaniv Erlich

## Abstract

Short Tandem Repeats are among the most polymorphic loci in the human genome. These loci play a role in the etiology of a range of genetic diseases and have been frequently utilized in forensics, population genetics, and genetic genealogy. Despite this plethora of applications, little is known about the variation of most STRs in the human population. Here, we report the largest-scale analysis of human STR variation to date. We collected information for nearly 700,000 STR loci across over 1,000 individuals in phase 1 of the 1000 Genomes Project. This process nearly saturated common STR variations. After employing a series of quality controls, we utilize this call set to analyze determinants of STR variation, assess the human reference genome’s representation of STR alleles, find STR loci with common loss-of-function alleles, and obtain initial estimates of the linkage disequilibrium between STRs and common SNPs. Overall, these analyses further elucidate the scale of genetic variation beyond classical point mutations. The resource is publicly available at http://strcat.teamerlich.org/ both in raw format and via a graphical interface.

## Introduction

STRs are abundant repetitive elements that are comprised of recurring DNA motifs of 2-6 bases. These loci are highly prone to mutations due to their susceptibility to slippage events during DNA replication (Ellegren 2004). To date, STR mutations have been linked to at least 40 monogenic disorders (Pearson et al. 2005; Mirkin 2007), including a range of neurological conditions such as Huntington’s disease, amyotrophic lateral sclerosis, and certain types of ataxia. Some disorders, such as Huntington’s disease, are triggered by the expansion of a large number of repeat units. In other cases, such as oculopharyngeal muscular dystrophy, the pathogenic allele is only two repeat units away from the wild-type allele (Brais et al. 1998; Amiel et al. 2004). In addition to Mendelian conditions, multiple studies in model organisms have suggested that STR variations contribute to an array of complex traits (Gemayel et al. 2010)], ranging from the period of the circadian clock in *Drosophila* (Sawyer et al. 1997) to gene expression in yeast (Vinces et al. 2009).

Beyond their importance to medical genetics, STR variations convey high information content due to their rapid mutations and multi-allelic spectra. Population genetics studies have utilized STRs in a wide-range of methods to find signatures of selection and to elucidate mutation patterns in nearby SNPs (Tishkoff et al. 2001; Sun et al. 2012). In DNA forensics, STRs play a significant role. Both the US and the European forensic DNA databases solely rely on these loci to create genetic fingerprints (Kayser and de Knijff 2011). Finally, the vibrant genetic genealogy community extensively uses these loci to develop impressive databases containing lineages for hundreds of thousands of individuals (Khan and Mittelman 2013).

Despite their utility, systematic data about the landscape of STR variations in the human population is far from comprehensive. Currently, most of the genetic information concerns a few thousand loci that were part of historical STR linkage and association panels in the pre SNP-array era (Broman et al. 1998; Tamiya et al. 2005) and several hundred loci involved in forensic analysis, genetic genealogy, or genetic diseases (Ruitberg et al. 2001; Pearson et al. 2005). In total, there are only 5,500 loci under the microsatellite category in dbSNP139. For the vast majority of STR loci, little is known about their normal allelic ranges, frequency spectra, and population differences. This knowledge gap largely stems from the absence of high-throughput genotyping techniques for these loci (Jorgenson and Witte 2007). Capillary electrophoresis offers the most reliable method to probe these loci, but this technology scales poorly. More recently, several studies have begun to explore STR loci with whole-genome sequencing datasets obtained from long read platforms such as Sanger sequencing (Payseur et al. 2011) and 454 (Molla et al. 2009; Duitama et al. 2014). However, due to the relatively low throughput of these platforms, these studies analyzed STR variations in only a few genomes.

Illumina sequencing has the potential to profile STR variations on a population-scale. However, STR variations present significant challenges for standard sequence analysis frameworks (Treangen and Salzberg 2012). In order to reduce computation time, most alignment algorithms employ heuristics that reduce their tolerance to align reads with large indels, hampering alignment of STRs with large contractions or expansions. In addition, due to the repetitive nature of STRs, the PCR steps involved in sample preparation induce *in vitro* slippage events (Hauge and Litt 1993). These events, called stutter noise, generate erroneous reads that mask the true genotypes. Because of these issues, previous large-scale efforts to catalog genetic variations have omitted STRs from their analyses (The 1000 Genomes Consortium 2012; Tennessen et al. 2012; Montgomery et al. 2013) and early attempts to analyze STRs using the 1000 Genomes data were mainly focused on exonic regions (McIver et al. 2013) or extremely short STRs regions with a relatively small number of individuals based on the native indel callset (Ananda et al. 2013).

In our previous studies, we created publicly available programs that specialize in STR profiling using Illumina whole-genome sequencing data (Gymrek et al. 2012; Highnam et al. 2013). Recently, we deployed one of these tools (lobSTR) to accurately genotype STRs on the Y chromosome of anonymous individuals in the 1000 Genomes Project to infer their surnames (Gymrek et al. 2013), demonstrating the potential utility of STR analysis from Illumina sequencing.

Here, we performed a genome-wide analysis of STR variation in the human population. First, we developed a quantitative approach to define plausible STR loci in the human reference genome. Next, we characterized STR variations in these loci in individuals from the 1000 Genomes Project (2012). We then used an array of external and internal controls to examine the quality of the STR variations. We found that while individual genotype calls were noisy, our catalog provided relatively accurate summary statistics for most of the loci, such as heterozygosity and the typical allelic spectrum. Finally, we examined the utility of our call set for applications in population genetics and personalized medicine.

The full catalog of STR variations is publicly available on http://strcat.teamerlich.org in VCF format. In addition, the website provides a series of graphical interfaces to search STR loci with specific biological properties, obtain summary statistics such as allelic spectrum and heterozygosity rates, and view the supporting raw sequencing reads.

## Results

### Identifying STR loci in the human genome

The first task in creating a catalog of STR variations is to determine the loci in the human reference that should be considered as STRs. This problem primarily stems from the lack of consensus in the literature as to how many copies of a repeat distinguish an STR from genomic background (Leclercq et al. 2007; Fondon et al. 2012; Schaper et al. 2012). For example, is (AC)_2_ an STR? What about (AC)_3_ or (AC)_10_? Furthermore, as sporadic bases can interrupt repetitive DNA sequences, purity must also be taken into account when deciding whether a locus is a true STR.

We employed a quantitative approach to identify STR loci in the reference genome. Multiple lines of study have proposed that the birth of an STR is a relatively rare event with complex biology (Ellegren 2004; Buschiazzo and Gemmell 2006; Oliveira et al. 2006; Gemayel et al. 2010; Kelkar et al. 2011; Ananda et al. 2013). The transition from a proto-STR to a mature STR requires non-trivial mutations such as the arrival of a transposable element, slippage-induced expansion of the proto-STR, or precise point mutations that destroy non-repetitive gaps between two short repeat stretches. Based on these observations, it was suggested that randomly-shuffled DNA sequence should rarely produce mature STR sequences and therefore can be used as negative controls for STR discovery algorithms (Gemayel et al. 2010; Schaper et al. 2012). We utilized this approach to identify STR loci in the human genome while controlling the false positive rate (Supplemental Figure 1; **Supplemental Methods**). We first integrated the purity, composition, and length of putative STRs in the genome into a score using Tandem Repeats Finder [TRF] (Benson 1999). Then, we generated random DNA sequences using a second-order Markov chain with similar properties to the human genome (i.e. nucleotide composition and transition frequencies). We tuned the TRF score threshold such that only 1% of the identified STR loci in our collection were expected to be false positives. We then evaluated the false negative rate of our catalog using two methods (**Supplemental Methods**). First, we collected a preliminary call set of repeat number variability across the human population with a highly permissive definition of STR loci. We found that our catalog misses only ∼1% of loci that exhibited repeat variability in the permissive call set (Supplemental Table 1). Second, we also collated a set of about 850 annotated *bona-fide* STR loci, mainly from the CODIS forensic panel and Marshfield linkage panel. Only 12 (1.4%) of these markers were not included in the catalog based on the TRF score threshold. The results of the two validation methods suggest that our catalog includes ∼99% of the true STRs in the genome and has a false positive rate of about 1%.

Overall, our STR catalog includes approximately 700,000 loci in the human genome. About 75% of these loci are di and tetra-nucleotide STRs, while the remaining loci are tri, penta and hexa-nucleotide STRs (Supplemental Table 2). Approximately 4,500 loci overlap coding regions, of which ∼80% have either trimeric or hexameric repeat units.

### Profiling STRs in 1000 Genomes samples

We collected variations in these 700,000 STR loci from 1,009 individuals from phase 1 of the 1000 Genomes Project (**Methods**; Supplemental Table 3). These samples span populations from five continents and were subject to low coverage (∼5x) whole-genome sequencing. In addition, high coverage exome sequencing data was available for 975 of these samples and was integrated with the whole-genome raw sequencing files.

We used two distinct STR genotyping pipelines designed to analyze high-throughput sequencing data, namely lobSTR (Gymrek et al. 2012) and RepeatSeq (Highnam et al. 2013). The STR genotypes of the two tools were quite concordant and matched in 133,375,900 (93%) out of the 143,428,544 calls that were reported by both tools. We tested multiple methods to unify the two call sets in order to further improve the quality (Supplemental Figure 2; **Supplemental Methods**). However, all of these integration methods did not improve the accuracy. Since the lobSTR calls showed better quality for highly polymorphic STRs, we proceeded with the analysis of STR variations solely based on this call set.

On average, we collected STR genotypes for approximately 530 individuals per locus (Figure 1a) and 350,000 STR loci per individual (Figure 1b), accumulating a total of about 350 million STR genotypes in the catalog. We examined the marginal increase in the number of covered STR loci as a function of sample size (**Methods**; Figure 1c). Our analysis shows that after analyzing 100 samples, there is a negligible increase in the number of genotyped STRs. However, even with all of the data, 3% of STR loci are persistently absent from the catalog. The average reference allele length of the missing STR loci was 182bp compared to 31bp for the rest of the reference, suggesting that the missing STR loci have allele lengths beyond the read length of Illumina sequencing. We also examined the marginal increase of polymorphic STR loci with minor allele frequencies (MAF) greater than 1%. Again, we noticed an asymptote at 100 samples that reaches approximately 300,000 loci. These saturation analyses suggest that with the current sample size, the STR variation catalog virtually exhausted all loci with MAF>1% that can be observed with 100bp Illumina reads and our analysis pipeline.

**Figure 1:**
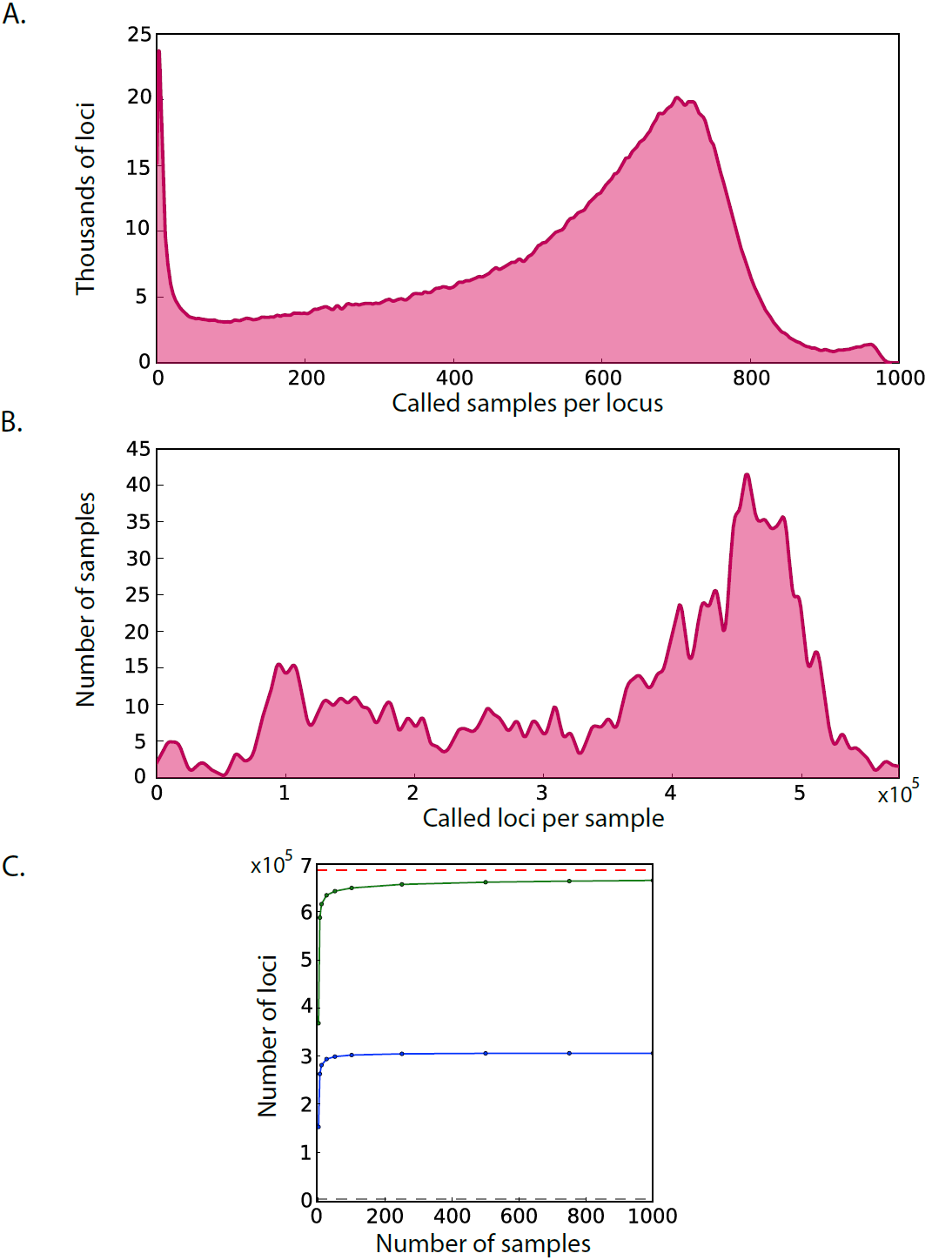
Call set statistics (A) Distribution of the number of called samples per locus. The average is 528 samples per STR with a standard deviation of 231 **(B) Distribution of the number of called loci per sample**. The average is 349,892 STR per sample with a standard deviation of 145,135 **(C) Saturation curves of the catalog**. The number of called loci (green) rapidly approaches the total number of STRs in the genome (red line). The number of called loci with a MAF>1% (blue) saturates after 100 samples and far exceeds the number of STR variants in dbSNP (grey line close to the X-axis).

### Quality assessment of STR loci

To initially assess the accuracy of our STR calls, we first examined patterns of Mendelian inheritance (MI) of STR alleles for three low-coverage trios present in the 1000 Genomes sample set. In total, we accumulated half a million genotypes calls. Without any read depth threshold, 94% of the STR loci followed MI (Figure 2a). The MI rates increased monotonically with read depth and restricting the analysis to loci with ≥10 reads increased the Mendelian inheritance to over 97%.

**Figure 2:**
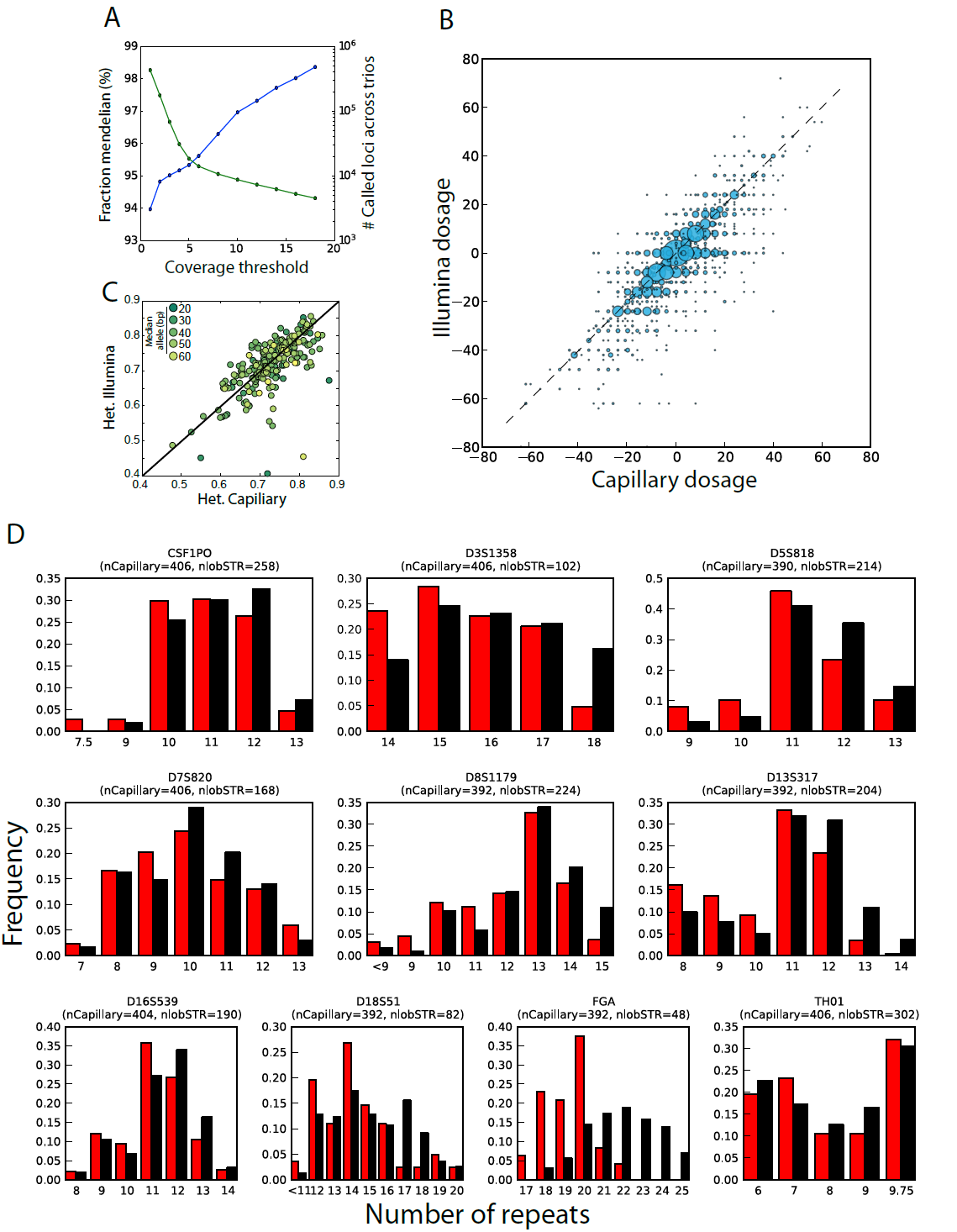
Quality assessments of the STR catalog (A) Consistency of lobSTR calls with Mendelian inheritance. The blue line denotes the fraction of STR loci that followed Mendelian inheritance as a function of the read coverage threshold. The green line denotes the total number of calls in the three trios that passed the coverage threshold **(B) Concordance between lobSTR and capillary electrophoresis genotypes**. The STR calls were taken from the highly polymorphic Marshfield panel. The dosage is reported as the sum of base pair differences from the NCBI reference. The area of each bubble is proportional to the number of calls of the dosage combination and the broken line indicates the diagonal. (**C**) **Comparison of heterozygosity rates for Marshfield panel STRs.** Color denotes the length of the median allele of the STR (dark-short; bright-long) **(D) Allelic spectra of the CODIS markers**. Red: lobSTR, black: capillary electrophoresis. nlobSTR and nCapillary indicate the number of allele called in the respective call sets.

Next, we compared the concordance of the calls in our catalog to those obtained using capillary electrophoresis, the gold standard for STR calling (**Methods**). We focused on datasets containing Marshfield and PowerPlex Y chromosome panel genotypes that are available for a subset of the 1000 Genomes individuals. These panels ascertain some of the most polymorphic STR loci, testing our pipeline in a challenging scenario. The Marshfield capillary panel (Rosenberg et al. 2005) reported 5,164 genotypes that overlapped with the lobSTR calls. These genotypes covered 157 autosomal STRs and were obtained from 140 individuals. The PowerPlex capillary panel reported 784 genotypes that overlapped with the lobSTR calls. The genotypes covered 17 Y-STRs and were obtained from 228 individuals. For each genotype in the two panels, we converted the reported alleles to STR dosages by summing the number of repeats after subtracting the reference allele. For example, if the genotype was 16bp/18bp and the reference allele was 14bp, the dosage of the locus was set to 2+4=6. For hemizygous loci, we just reported the difference from the reference allele. After regressing the lobSTR dosages with the capillary dosages, the resulting goodness of fit estimators (R^2^) were 0.71 for the autosomal genotypes and 0.94 for the Y chromosome genotypes (Figure 2b; Supplemental Figure 3). By further stratifying the autosomal calls by the capillary genotype, we found that lobSTR correctly reported 89.5% of all homozygous loci and recovered one or more alleles for 91.5% of all heterozygous loci, but only correctly reported both alleles for 12.8% of all heterozygous loci (Supplemental Table 4). For the Y chromosome, 95% of the lobSTR genotypes exactly matched the capillary genotypes for the PowerPlex Y panel (Supplemental Table 5).

Collectively, these results suggest that allelic dropouts are the primary source of noise in the call set but that the individual allele lengths are relatively accurate. This statement is supported by the relatively good accuracy of the hemizygous Y chromosome call set and the alleles in the homozygous loci. In general, allelic dropouts are quite expected given the relatively low sequencing coverage but are also known to be an issue in genotyping STRs with capillary electrophoresis (Pompanon et al. 2005).

We performed various analyses that demonstrate that allelic dropouts do not hamper the ability to deduce population-scale patterns of human STR variation. First, we examined the heterozygosity rates of the Marshfield STRs in three European subpopulations in our call set (CEU, GBR and FIN). We found that the heterozygosity rates were significantly correlated (R=0.68; p<10^-30^) between the lobSTR and the capillary results (Figure 2c). In addition, we analyzed the allelic spectra of European populations for over 200 STR loci in the Marshfield panel (Supplemental Figure 4). We found that in most cases, lobSTR and the capillary spectra matched in the median and interdecile range of the reported allelic lengths. We also inspected the allelic frequency spectra of STRs that are part of the forensic CODIS test panel using a similar procedure (Figure 2d). A previous study reported the spectra of these loci by capillary electrophoresis genotyping in ∼200 Caucasians in the United States (Budowle et al. 1999). Again, these comparisons resulted in similar patterns for eight of the ten analyzed markers. We found marked biases only for FGA and D18S51, with lobSTR reporting systematically shorter alleles. As the maximal allele sizes of these two loci are over 80bp, the long alleles are seldom spanned by Illumina reads, creating a bias toward shorter alleles. However, only 3-4% of the STRs in our catalog have a reference allele exceeding the lengths of these loci. Therefore, we do not expect this bias to affect the allelic spectra of most loci.

To further assess the utility of our catalog, we tested its ability to replicate known population genetics trends. We specifically wondered about the quality of the most variable STR loci in the catalog. One hypothesis is that these loci are just extreme cases of genotyping errors; an alternative hypothesis is that these loci are truly polymorphic and can provide useful observations about the underlying populations. We first compared the heterozygosities of the 10% most variable autosomal loci across ten different subpopulations from Africa, East Asia, and Europe. Consistent with the Out-of-Africa bottleneck (Stoneking and Krause 2011), we found that the genetic diversity of the African subpopulations significantly exceeded those of Europe and East Asia (sign test; p < 10^-50^ for any African non-African pair) (Figure 3a; Supplemental Table 6). Second, we focused on the 100 most heterozygous autosomal loci in our catalog and inspected the ability of STRUCTURE (Pritchard et al. 2000) to cluster a subset of the samples into three main ancestries in an unsupervised manner. Our results show that all of these samples clustered distinctly by geographical region (Figure 3b). These analyses demonstrate that even the most variable loci in the catalog still convey valid genetic information that can be useful for population genetic analyses.

**Figure 3:**
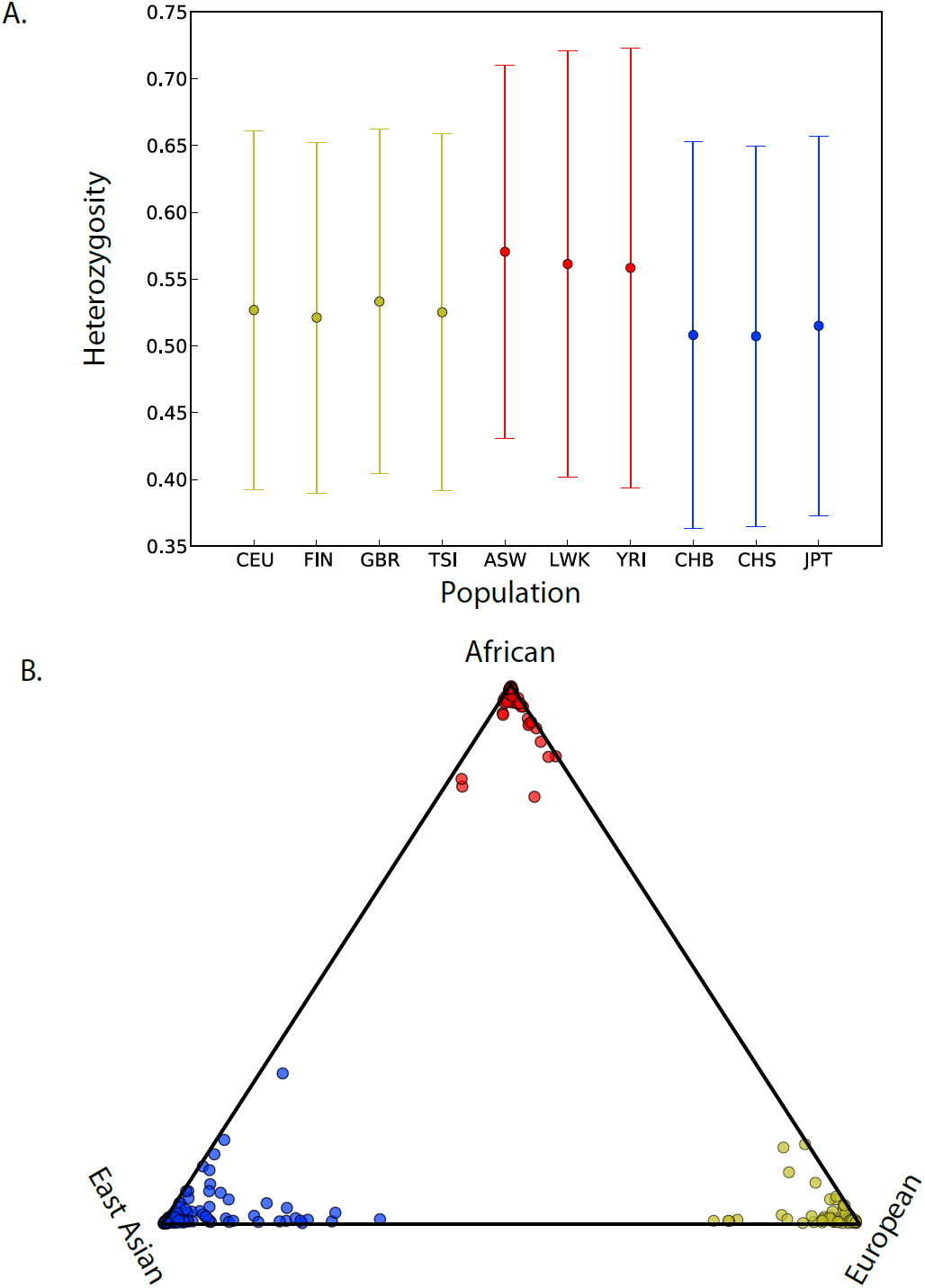
Evaluation of the STR catalog for population genetics (A) Genetic diversity of the 10% most heterozygous autosomal loci in different populations of the 1000 Genomes. Yellow: European, Red: African, Blue: East Asian. The mean heterozygosity (dot) of the African subpopulations consistently exceeds those of the non-African subpopulations. The whiskers extend to ± one standard deviation. See Supplemental Table 3 for population abbreviations **(B) STRUCTURE clustering based on the 100 most polymorphic autosomal STR loci.** Each subpopulation clusters tightly by geographic origin. Color labels as in (A).

In summary, the multiple lines of quality assessment suggest that our catalog can be used to infer patterns of human STR variations such as heterozygosity, allelic spectra, and population structure. The most notable shortcoming of the catalog is allelic dropouts stemming from the low sequencing coverage of the 1000 Genomes. However, the experiments above suggest that valuable summary statistics can be extracted from the call set despite this caveat.

### Patterns of STR variation

Next, we analyze the sequence determinants of STR variation. We found that for noncoding STRs, variability monotonically decreased with motif length (Figure 4a). In contrast, loci with trimeric and hexameric motifs were the most polymorphic among coding STRs. These STR loci can vary without introducing frameshift mutations and therefore may be exposed to weaker purifying selection. In addition, coding STRs demonstrated significantly reduced heterozygosity compared to noncoding STRs for periods 2-5bp (Mann-Whitney U test; p < 0.01, Supplemental Table 7). Hexameric STRs, on the other hand, showed no statistically significant difference in variability between loci in these two classes. In addition to period, we also found that the length of an STR significantly affects its variability (Supplemental Table 8). For pure STRs without any interruptions, we found a nearly monotonic increase in the variability of loci as the length of the most common allele increased (Figure 4b). Similar trends also applied for STRs with various levels of impurities, albeit with a reduced magnitude of effect for less pure loci (Supplemental Figure 5). These patterns of STR variability agree with previous small-scale studies of STRs genotyped with capillary electrophoresis (Pemberton et al. 2009) and match patterns of STR variation observed in a single deeply sequenced genome (Gymrek et al. 2012), providing additional support for the validity of our call set.

**Figure 4:**
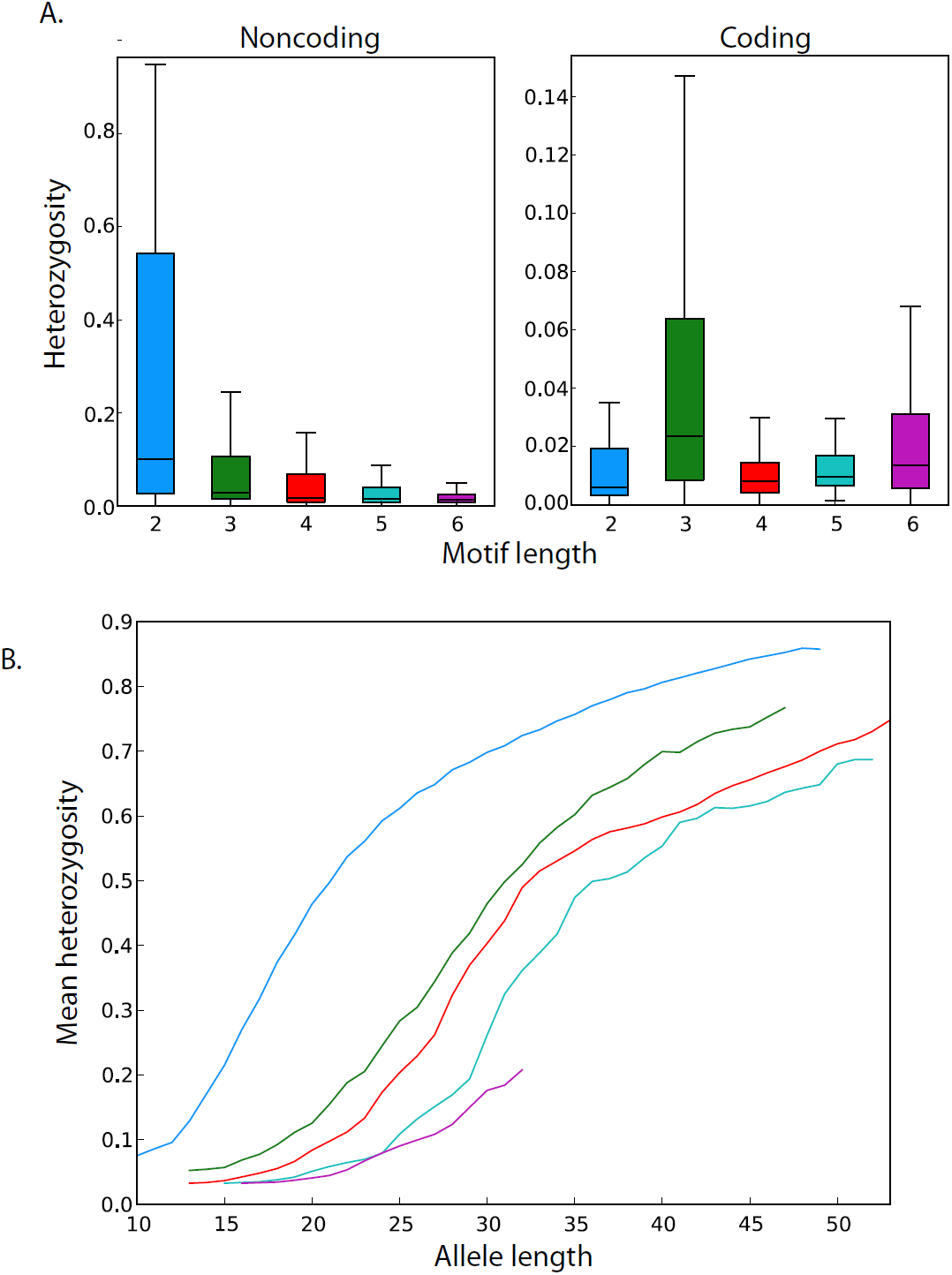
Determinants of STR variability (A) Motif length and coding capabilities as determinants of STR variability. STR heterozygosity monotonically decreases with period for noncoding loci and is generally reduced in non-coding (left) versus coding regions (right). The box extends from the lower to upper quartiles of the heterozygosity distribution and the interior line indicates the median. The whiskers extend to the most extreme points within 1.5*IQR of the quartiles. **(B) STR variability as a function of the wild-type allele (base pairs).** The mean heterozygosity of STRs increases monotonically with allele length for each of the five STR periods. Analysis was restricted to STRs with wild-type allele matching the reference and with no indels or SNPs interrupting the STR motif (blue: 2mer, green: 3mer, red: 4mer, cyan: 5mer, purple: 6mer).

We also wondered about the prototypical pattern of variation of an STR locus in terms of the number of alleles and their distribution. We found that 30% of STRs have a common polymorphism with at least two alleles with frequencies above 5%. Dinucleotide STRs have the highest rate, with 48% of these loci displaying a common polymorphism. Moreover, 30% of all dinucleotide STRs have more than 3 alleles with a frequency above 5%. On the other hand, hexanucleotide STRs have the lowest common polymorphism rate, with only 13% of these loci displaying a common polymorphism (Supplemental Figure 6a; Supplemental Table 9). Next, we turned to find the prototypal allelic spectra of STR variations. For each STR, we normalized the reported alleles such that they reflected the distance in number of repeats from the wild-type allele in the locus. Then, we generated histograms that show the allelic spectra by aggregating all the alleles of STRs with the same motif length. This coarse-grained picture was similar across repeat lengths (Supplemental Figure 6b). The allelic spectrum of an STR is unimodal and relatively symmetric. There is one, highly prevalent wild-type allele, two less common alleles with one repeat above and below the wild-type allele, and a range of rare alleles with monotonic decreased frequency that reach over ±5 repeats from the wild-type allele.

### STRs in the NCBI reference and LoF analysis

We were interested in assessing how well the wild-type alleles are represented in the NCBI reference (Figure 5a). We found that for over 69,000 loci (10% of our reference set), the wild-type allele across the 1000 Genomes populations was at least one repeat away from the NCBI hg19 reference allele. In addition, 15,581 loci (2.25%) in the reference genome were 10bp or more away from the most common allele in our dataset. For STRs in coding regions, 48 loci (1.1% of coding STRs) had a wild-type allele that did not match the NCBI reference (Supplemental Table 10). In 46 out of 48 of the cases, these differences occurred for loci with trinucleotide or hexanucleotide repeats and conserved the reading frame. Moreover, the two wild-type frame-shifted alleles are unlikely to trigger the non-sense mediated decay pathway. The deletion of one 4bp unit in *DCHS2* occurs a few nucleotides before the annotated RefSeq stop codon. This variation slightly alters the location of the stop codon and affects only five amino acids in the C-terminus of the protein. The 14bp deletion in *ANKLE1* occurs in the last exon of the gene and introduces about 20 new amino acids to the tail of the protein.

**Figure 5:**
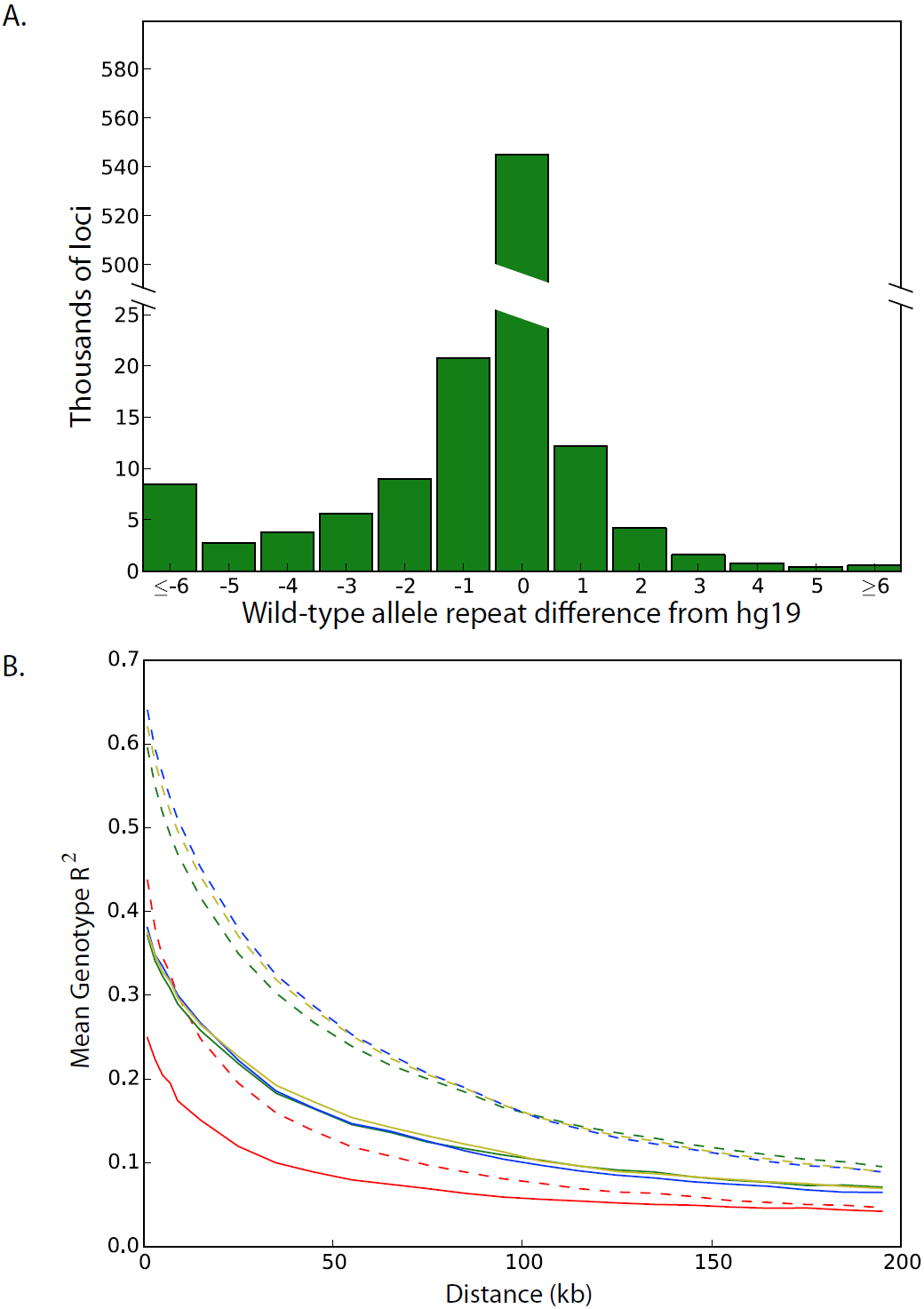
Population-scale analyses of STR variation (A) Distribution of base-pair differences between each locus’ wild-type and NCBI reference allele (B) Patterns of linkage disequilibrium for SNPs and STRs on the X chromosome. SNP-SNP LD (dashed lines) generally exceeds SNP-STR LD (solid lines) across a range of distances and for Africans (red), Admixed Americans (green), Europeans (yellow) and East Asians (blue).

We also sought to identify a confident set of STR loci with relatively common loss of function (LoF) alleles. To accomplish this goal, we considered only alleles supported by at least two reads and 30% of the total reads per called genotype. We further required that alleles be carried by 10 or more samples. Seven common LoF alleles across five genes passed this criterion: *DCHS2*, *FAM166B*, *GP6*, *SLC9A8*, and *TMEM254* (Supplemental Table 11). Out of these 5 genes, only *GP6* has known implications for a Mendelian condition: a mild platelet-type bleeding disorder (Dumont et al. 2009; Hermans et al. 2009). However, the LoF mutation in this gene resides in the last exon and is unlikely to induce the non-sense mediated decay pathway. In conclusion, the LoF analysis indicates that common STR polymorphisms rarely disrupt the reading frame.

### Linkage disequilibrium between STRs and SNPs

The linkage disequilibrium (LD) structure of STRs and SNPs is largely unknown. On top of recombination events, the SNP-STR LD structure also absorbs STR back mutations that could further shift these pairs of loci towards equilibrium. However, there is minimal empirical data in the literature about the pattern of this LD structure, most of which pertains to a few hundred autosomal Marshfield markers (Payseur et al. 2008). To get a chromosome-wide estimate, we inspected STR loci on the hemizygous X chromosomes in male samples. Similar to the Y chromosome data, these calls do not suffer from allelic dropouts and are already phased with SNP alleles, conferring a technically reliable dataset for a chromosome-wide analysis.

We determined the LD in terms of the r^2^ between SNPs and STRs as a function of the distance between these markers. Only STRs and SNPs with common polymorphisms were used for the analysis. Hexameric STRs were not included due to the small sample size of 24 sites; for the other repeat motifs, we obtained hundreds to thousands of polymorphic markers. We stratified the STR-SNP LD based on the four major continental populations (Africa, Asia, Europe, and America) and contrasted them to the patterns for classical SNP-SNP LD (Figure 5b). In all cases, the SNP-SNP LD consistently exceeded mean STR-SNP LD. In addition, the African population demonstrated markedly reduced levels of SNP-STR LD and SNP-SNP LD, consistent with its larger effective population size. In general, dinucleotide STRs showed the weakest LD with nearby SNPs, which likely stems from their higher mutation rates (Supplemental Figure 7).

Overall, this analysis shows that the average SNP-STR LD is approximately half of the SNP-SNP LD of variations with the same distance on the X chromosome. Since the effective population size of the X chromosome is smaller than that of the autosome, the STR-SNP LD should be even smaller on the autosome. These results suggest that association studies with tagging SNPs might be considerably underpowered to detect loci with causal STRs, specifically dinucleotide loci.

## Discussion

In the last few years, population-scale sequencing projects have made tremendous progress in documenting genetic variation across human populations. The 1000 Genomes Project has already reported approximately 40 million SNPs, 1.4 million insertion and deletions, and over 10,000 structural variants (1000 Genomes Consortium, 2012). Similar catalogs, albeit to lesser degrees of completeness, have been produced for other types of variations, such as LINE-1 insertions (Ewing and Kazazian 2011) and Alu repeat variations (Hormozdiari et al. 2011). Here, we presented a population-scale analysis of STR variation, adding another layer of genetic variation to existing catalogs.

Our analysis significantly augments the level of knowledge of STR variations. Initially, dbSNP reported data for only 5,500 STR loci. Our catalog provides data on close to 700,000 STR loci, which encompasses 97% of the STRs in the genome, and contains over 300,000 STR loci with a MAF of over 1%. One caveat of our catalog is the low reliability of individual genotypes due to allelic dropout. Nonetheless, we showed using multiple lines of analysis that summary statistic results such as frequency spectra and variation trends can be extracted from the catalog.

The landscape of STR variations in the apparently healthy 1000 Genomes individuals suggests several rules of thumbs for analyzing STR variations for medical sequencing. Previous work found that membrane proteins of several pathogens contain STR loci with non-triplet motifs whose variations can be beneficial to the organism (Gemayel et al. 2010). These STRs confer high evolvability and adaptability of these proteins by dynamically changing the reading frame. In contrast, our data suggests that for the vast majority of human proteins, frame-shift mutations in their STR regions are not favorable. Only a handful of STR loci harbor common frame-shift polymorphisms and half of the LoF alleles create a very small change in the C-terminus tail of the protein. Based on these observations, we hypothesize that most of the non-triplet coding STRs are not-well tolerated and are exposed to negative selection similarly to a regular indels in the same region. Therefore, it is advisable for medical sequencing projects to analyze these loci as well and treat them as regular LoF alleles rather than filtering them. This rule of thumb is well-echoed in a recent study of medullary cystic kidney disease type 1 that implicated the genetic pathology in a frame-shift mutation caused by a length change of a homopolymer run (Kirby et al. 2013). For in-frame STR variations, our call set contains deep allelic spectra of most of these loci, providing reference distributions of apparently healthy alleles. These spectra can be used to identify atypical STR alleles and might serve as an indicator for pathogenicity.

Another open question raised from our findings is the potential contribution of STRs to complex traits. Using the prototypical allelic spectra, we estimate that the average additive variances are 3, 0.7, 0.4, 0.25 and 0.1 for 2-6mer STRs, respectively. Interestingly, the theoretical maximum of additive variance for a bi-allelic SNP dosage is 0.5, six times smaller than the observed variance of dinucleotide STRs. From a theoretical statistical genetics perspective, this suggests that causal dinucleotide STR loci could explain a considerable fraction of phenotypic variance even with a relatively modest effect size. Therefore, if each STR allele in a locus slightly changes a quantitative trait in a gradual manner, the net effect on the phenotypic variance could be quite large due to the wide range of these alleles and their relatively high frequencies. Interestingly, we found that dinucleotide loci show relatively weak LD with SNPs. This complicates finding these STRs in association studies with SNP arrays. With the theoretical potential of STRs to contribute to phenotypic variance on one hand and their weaker LD to tagging SNPs on the other hand, there is a possibility that these loci contribute to the missing heritability phenomenon of complex traits (Manolio et al. 2009). STR variations are yet to be systematically analyzed by GWAS association studies and our hope is that this catalog can be a reference point for future analyses.

## Methods

### Call set generation

The raw sequencing files of Phase 1 of the 1000 Genomes Project were analyzed.

The lobSTR calls were generated using computing resources hosted by Amazon Web Services, GitHub version 8a6aeb9 of the lobSTR genotyper and Github version a85bb7f of the lobSTR allelotyper (https://github.com/mgymrek/lobstr-code). In particular, the lobSTR genotyper was run using the options fft-window-size=16, fft-window-step=4 and bwaq=15. PCR duplicates were removed from the resulting BAM files for each experiment using *samtools*. The individual BAMs were merged by population and the lobSTR allelotyper was run using all population BAMs concurrently, the include-flank option and version 2.0.3 of lobSTR’s Illumina PCR stutter model.

RepeatSeq (available http://github.com/adaptivegenome/repeatseq) was run using default parameters on the read alignments produced by the 1000 Genomes project.

For both programs, we used the 700,000 STR list that were constructed by the second-order Markov framework (**Supplemental Methods**).

### Estimating the numbers of samples per locus and loci per sample

The distributions of the call set parameters were smoothed using the gaussian_kde function in the *scipy.stats* python package. Covariance factors of 0.01 and 0.025 were used to smooth the samples per locus and loci per sample distributions, respectively.

### Saturation analysis

We determined the number of loci with calls for sample subsets containing 1, 5, 10, 25, 50, 100, 250, 500, 750 and 1000 individuals. In particular, we began by randomly selecting 1 individual. To create a subset of 5 individuals, we then added 4 more random individuals and son on. For each of these sample subsets, we determined the number of loci with one or more STR calls across all samples in the subset. We repeated this whole process 10 times and used the median number of called loci across each of the 10 repetitions to create the saturation profile for all loci.

We also determined whether loci had a MAF > 1% using all 1009 samples. We then used a procedure analogous to the one described above to select subsets of samples and determine whether or not each of these loci had a corresponding call in each subset. This procedure resulted in the saturation profile for loci with MAF > 1%.

### Mendelian inheritance

The three low-coverage trios contained within the dataset consisted of the following sample sets: HG00656, HG00657, HG00702 (trio 1), NA19661, NA19660, NA19685 (trio 2) and NA19679, NA19678, NA19675 (trio 3). To assess the consistency with Mendelian inheritance for a given trio, only loci for which all three samples had calls were analyzed. The coverage assigned to each trio of calls corresponded to the minimum coverage across the three samples.

### Capillary electrophoresis comparison

Marshfield genotypes (Rosenberg et al. 2005) were downloaded from http://www.stanford.edu/group/rosenberglab/data/rosenbergEtAl2005/combined-1048.stru. Prior to comparing genotypes, offsets were calculated to match the lobSTR calls to the length of the Marshfield PCR products. For each locus, all observed offsets were considered and scored and the optimally scoring offset across all samples was selected. In particular, for each sample, an offset was scored as a 1, 0.5, 0.25 or 0 if the lobSTR calls matched exactly, were homozygous and recovered one Marshfield allele, were heterozygous and recovered one Marshfield allele or did not match at all, respectively. Only loci with at least 20 calls were considered in the comparison. Finally, the Pearson correlation coefficient was calculated using the sum of the allele length differences from hg19 for each locus in each sample.

Y chromosome PowerPlex genotypes were downloaded from the 1000 Genomes Y chromosome working group FTP site. Offsets were once again calculated to match the length of the PCR products to the lobSTR calls. For each locus, the offset was calculated as the most common difference between the lobSTR and PowerPlex genotypes across samples. Only loci with at least 5 calls were considered in the comparison and the R^2^ was calculated between the allele length differences from hg19 for each locus in each sample. In addition, the 15 heterozygous lobSTR calls were ignored.

Slopes and R^2^ values for STR dosage comparisons were calculated using the linregress function in the scipy.stats package. To mitigate the effects of outliers, we explored using regular linear regression, regression with a zero intercept and L1 penalized regression. The resulting slopes were essentially invariant to the calculation method and so statistics were reported based on traditional linear regression.

### Summary statistic comparisons

The allelic spectra of the Marshfield panel were downloaded from http://research.marshfieldclinic.org/genetics/genotypingData_Statistics/markers/ and parsed using a custom Perl script (available upon request from the authors). Samples from the CEU, GBR, TSI, and FIN subpopulations were analyzed, and only markers with more than 50 calls were included.

We utilized all of the lobSTR calls for the CEU, GBR and FIN subpopulations to generate the lobSTR frequency spectra for each CODIS marker. Spectra were not available for 3 of the CODIS markers (D21S11, VWa, TPOX). D21S11 is too long to be spanned by Illumina reads; we had annotation difficulties for VWa and TPOX (assigning the correct STR in hg19 to the NIST STR). We then compared the available frequency spectra to those published for a Caucasian population in the United States (Budowle et al. 1999).

Because of some annotation differences between the capillary data and our reference locations, we shifted the lobSTR spectra for the D8S1179 marker by +2 repeat units. Finally, repeat lengths for which the maximum frequency was less than 2% were not displayed.

### Comparison of population heterozygosity

To obtain accurate measures of heterozygosity, autosomal STR loci with less than 30 calls in any of the 10 subpopulations considered were ignored. Of the remaining loci, the 10% most heterozygous (24,637 loci) were selected and their means and standard deviations were calculated using the mean and stdev functions in the python numpy package. To determine whether a pair of populations had systematically different heterozygosity at these loci, we paired the heterozygosities for each locus and counted the number of pairs in which population A had a larger heterozygosity than population B. Ignoring the relatively small number of loci in which heterozygosities were identical, the p-value for this over/underrepresentation was then calculated using the cdf function in the scipy.stats.binom python package.

### Sample clustering

*STRUCTURE* version 2.3.4 was utilized to perform the MCMC-based clustering. The program was run using MAXPOPS=3, BURNIN=500000, NUMREPS=1000000, no prior population information, unphased genotypes, the admixture model and no linkage disequilibrium. All 321 samples from the JPT, CHB, YRI and CEU subpopulations present in the data were clustered based on the 100 most heterozygous autosomal STRs with at least 750 called samples. Samples for which at least 75% of the selected makers were missing calls were not including in the resulting visualization. The final triangle plot therefore contained data for 71, 80, 81, and 82 samples from the CEU, CHB, JPT and YRI populations, respectively.

### STR variability trends

Analysis was restricted to STRs with at least 100 called samples. STRs that overlapped an annotated RefSeq translated region were regarded as coding and these annotations were downloaded from the UCSC table browser on 2/11/2014. The mannwhitneyu function in the scipy.stats python package was used to test for significant differences between coding and non-coding STR heterozygosity. For analyses related to allele length, STRs were further restricted to those with wildtype matching the hg19 reference to enable calculation of the locus’ purity. In particular, the purity of each of these STRs was calculated as the fraction of possible positions within the STR region where the subsequent bases corresponded to a cyclic permutation of the STR’s motif. The *pearsonr* function in the *scipy.stats* python package was used to calculate the Pearson correlation coefficients and their associated p-values, where each STR’s length and heterozygosity represented an individual point. Finally, to generate the plots of heterozygosity vs. length, the heterozygosity for each length was calculated as the mean variability of loci within 2bp.

### Assessing linkage disequilibrium

In order to avoid phasing SNPs and STRs, we only analyzed X chromosome genotypes in male samples. SNP calls for the corresponding samples were obtained from the 1000 Genomes Phase 1 11/23/2010 release and any pseudoautosomal loci were ignored. Analysis of STR-SNP LD was restricted to STR loci with both a heterozygosity of at least 9.5% and at least 20 genotypes for each super population (African, East Asian, European and Ad Mixed American). For each STR that met this requirement, we identified all SNPs within 200 KB of the STR start coordinate. After filtering out SNPs with a MAF below 5% in any of the four super populations, we calculated the level of LD for the remaining STR-SNP pairs. In particular, the R^2^ was calculated between the SNP genotype indicator variable and the base pair difference of the STR from the reference.

For SNP-SNP LD calculations, a seed SNP was identified for each STR meeting the aforementioned requirements. In particular, the SNP closest to the STR’s start coordinate with MAF > 5% for each super population was selected. If no such SNP existed within 1Kb, no SNP was selected and the STR was omitted from the STR-SNP LD analysis. Otherwise, we identified all SNPs within 200 KB of the seed SNP and one again removed SNPs with a MAF < 5% in any of the super populations. The LD between the seed SNP and each of these remaining SNPs was then assessed as the R^2^ between the two SNP genotype indicator variables.

## Acknowledgements

M.G. is supported by the National Defense Science & Engineering Graduate Fellowship. Y.E. is an Andria and Paul Heafy Family Fellow and holds a Career Award at the Scientific Interface from the Burroughs Wellcome Fund. This study was funded by a gift from Cathy and Jim Stone and an AWS Education Grant award. The authors thank Chris Taylor Smith, Wei Wei, Qasim Ayub, and Yali Xue for providing the results of the Y-STR panel for the 1000 Genomes individuals and the 1000 Genomes Project members for useful discussions.

The members of the 1000 Genomes project are: Altshuler DM, Durbin RM, Abecasis GR, Bentley DR, Chakravarti A, Clark AG, Donnelly P, Eichler EE, Flicek P, Gabriel SB, Gibbs RA, Green ED, Hurles ME, Knoppers BM, Korbel JO, Lander ES, Lee C, Lehrach H, Mardis ER, Marth GT, McVean GA, Nickerson DA, Schmidt JP, Sherry ST, Wang J, Wilson RK, Gibbs RA, Dinh H, Kovar C, Lee S, Lewis L, Muzny D, Reid J, Wang M, Wang J, Fang X, Guo X, Jian M, Jiang H, Jin X, Li G, Li J, Li Y, Li Z, Liu X, Lu Y, Ma X, Su Z, Tai S, Tang M, Wang B, Wang G, Wu H, Wu R, Yin Y, Zhang W, Zhao J, Zhao M, Zheng X, Zhou Y, Lander ES, Altshuler DM, Gabriel SB, Gupta N, Flicek P, Clarke L, Leinonen R, Smith RE, Zheng-Bradley X, Bentley DR, Grocock R, Humphray S, James T, Kingsbury Z, Lehrach H, Sudbrak R, Albrecht MW, Amstislavskiy VS, Borodina TA, Lienhard M, Mertes F, Sultan M, Timmermann B, Yaspo ML, Sherry ST, McVean GA, Mardis ER, Wilson RK, Fulton L, Fulton R, Weinstock GM, Durbin RM, Balasubramaniam S, Burton J, Danecek P, Keane TM, Kolb-Kokocinski A, McCarthy S, Stalker J, Quail M, Schmidt JP, Davies CJ, Gollub J, Webster T, Wong B, Zhan Y, Auton A, Gibbs RA, Yu F, Bainbridge M, Challis D, Evani US, Lu J, Muzny D, Nagaswamy U, Reid J, Sabo A, Wang Y, Yu J, Wang J, Coin LJ, Fang L, Guo X, Jin X, Li G, Li Q, Li Y, Li Z, Lin H, Liu B, Luo R, Qin N, Shao H, Wang B, Xie Y, Ye C, Yu C, Zhang F, Zheng H, Zhu H, Marth GT, Garrison EP, Kural D, Lee WP, Leong WF, Ward AN, Wu J, Zhang M, Lee C, Griffin L, Hsieh CH, Mills RE, Shi X, von Grotthuss M, Zhang C, Daly MJ, DePristo MA, Altshuler DM, Banks E, Bhatia G, Carneiro MO, del Angel G, Gabriel SB, Genovese G, Gupta N, Handsaker RE, Hartl C, Lander ES, McCarroll SA, Nemesh JC, Poplin RE, Schaffner SF, Shakir K, Yoon SC, Lihm J, Makarov V, Jin H, Kim W, Kim KC, Korbel JO, Rausch T, Flicek P, Beal K, Clarke L, Cunningham F, Herrero J, McLaren WM, Ritchie GR, Smith RE, Zheng-Bradley X, Clark AG, Gottipati S, Keinan A, Rodriguez-Flores JL, Sabeti PC, Grossman SR, Tabrizi S, Tariyal R, Cooper DN, Ball EV, Stenson PD, Bentley DR, Barnes B, Bauer M, Cheetham R, Cox T, Eberle M, Humphray S, Kahn S, Murray L, Peden J, Shaw R, Ye K, Batzer MA, Konkel MK, Walker JA, MacArthur DG, Lek M, Sudbrak R, Amstislavskiy VS, Herwig R, Shriver MD, Bustamante CD, Byrnes JK, De La Vega FM, Gravel S, Kenny EE, Kidd JM, Lacroute P, Maples BK, Moreno-Estrada A, Zakharia F, Halperin E, Baran Y, Craig DW, Christoforides A, Homer N, Izatt T, Kurdoglu AA, Sinari SA, Squire K, Sherry ST, Xiao C, Sebat J, Bafna V, Ye K, Burchard EG, Hernandez RD, Gignoux CR, Haussler D, Katzman SJ, Kent WJ, Howie B, Ruiz-Linares A, Dermitzakis ET, Lappalainen T, Devine SE, Liu X, Maroo A, Tallon LJ, Rosenfeld JA, Michelson LP, Abecasis GR, Kang HM, Anderson P, Angius A, Bigham A, Blackwell T, Busonero F, Cucca F, Fuchsberger C, Jones C, Jun G, Li Y, Lyons R, Maschio A, Porcu E, Reinier F, Sanna S, Schlessinger D, Sidore C, Tan A, Trost MK, Awadalla P, Hodgkinson A, Lunter G, McVean GA, Marchini JL, Myers S, Churchhouse C, Delaneau O, Gupta-Hinch A, Iqbal Z, Mathieson I, Rimmer A, Xifara DK, Oleksyk TK, Fu Y, Liu X, Xiong M, Jorde L, Witherspoon D, Xing J, Eichler EE, Browning BL, Alkan C, Hajirasouliha I, Hormozdiari F, Ko A, Sudmant PH, Mardis ER, Chen K, Chinwalla A, Ding L, Dooling D, Koboldt DC, McLellan MD, Wallis JW, Wendl MC, Zhang Q, Durbin RM, Hurles ME, Tyler-Smith C, Albers CA, Ayub Q, Balasubramaniam S, Chen Y, Coffey AJ, Colonna V, Danecek P, Huang N, Jostins L, Keane TM, Li H, McCarthy S, Scally A, Stalker J, Walter K, Xue Y, Zhang Y, Gerstein MB, Abyzov A, Balasubramanian S, Chen J, Clarke D, Fu Y, Habegger L, Harmanci AO, Jin M, Khurana E, Mu XJ, Sisu C, Li Y, Luo R, Zhu H, Lee C, Griffin L, Hsieh CH, Mills RE, Shi X, von Grotthuss M, Zhang C, Marth GT, Garrison EP, Kural D, Lee WP, Ward AN, Wu J, Zhang M, McCarroll SA, Altshuler DM, Banks E, del Angel G, Genovese G, Handsaker RE, Hartl C, Nemesh JC, Shakir K, Yoon SC, Lihm J, Makarov V, Degenhardt J, Flicek P, Clarke L, Smith RE, Zheng-Bradley X, Korbel JO, Rausch T, Sttz AM, Bentley DR, Barnes B, Cheetham R, Eberle M, Humphray S, Kahn S, Murray L, Shaw R, Ye K, Batzer MA, Konkel MK, Walker JA, Lacroute P, Craig DW, Homer N, Church D, Xiao C, Sebat J, Bafna V, Michaelson JJ, Ye K, Devine SE, Liu X, Maroo A, Tallon LJ, Lunter G, Iqbal Z, Witherspoon D, Xing J, Eichler EE, Alkan C, Hajirasouliha I, Hormozdiari F, Ko A, Sudmant PH, Chen K, Chinwalla A, Ding L, McLellan MD, Wallis JW, Hurles ME, Blackburne B, Li H, Lindsay SJ, Ning Z, Scally A, Walter K, Zhang Y, Gerstein MB, Abyzov A, Chen J, Clarke D, Khurana E, Mu XJ, Sisu C, Gibbs RA, Yu F, Bainbridge M, Challis D, Evani US, Kovar C, Lewis L, Lu J, Muzny D, Nagaswamy U, Reid J, Sabo A, Yu J, Guo X, Li Y, Wu R, Marth GT, Garrison EP, Leong WF, Ward AN, del Angel G, DePristo MA, Gabriel SB, Gupta N, Hartl C, Poplin RE, Clark AG, Rodriguez-Flores JL, Flicek P, Clarke L, Smith RE, Zheng-Bradley X, MacArthur DG, Bustamante CD, Gravel S, Craig DW, Christoforides A, Homer N, Izatt T, Sherry ST, Xiao C, Dermitzakis ET, Abecasis GR, Kang HM, McVean GA, Mardis ER, Dooling D, Fulton L, Fulton R, Koboldt DC, Durbin RM, Balasubramaniam S, Keane TM, McCarthy S, Stalker J, Gerstein MB, Balasubramanian S, Habegger L, Garrison EP, Gibbs RA, Bainbridge M, Muzny D, Yu F, Yu J, del Angel G, Handsaker RE, Makarov V, Rodriguez-Flores JL, Jin H, Kim W, Kim KC, Flicek P, Beal K, Clarke L, Cunningham F, Herrero J, McLaren WM, Ritchie GR, Zheng-Bradley X, Tabrizi S, MacArthur DG, Lek M, Bustamante CD, De La Vega FM, Craig DW, Kurdoglu AA, Lappalainen T, Rosenfeld JA, Michelson LP, Awadalla P, Hodgkinson A, McVean GA, Chen K, Tyler-Smith C, Chen Y, Colonna V, Frankish A, Harrow J, Xue Y, Gerstein MB, Abyzov A, Balasubramanian S, Chen J, Clarke D, Fu Y, Harmanci AO, Jin M, Khurana E, Mu XJ, Sisu C, Gibbs RA, Fowler G, Hale W, Kalra D, Kovar C, Muzny D, Reid J, Wang J, Guo X, Li G, Li Y, Zheng X, Altshuler DM, Flicek P, Clarke L, Barker J, Kelman G, Kulesha E, Leinonen R, McLaren WM, Radhakrishnan R, Roa A, Smirnov D, Smith RE, Streeter I, Toneva I, Vaughan B, Zheng-Bradley X, Bentley DR, Cox T, Humphray S, Kahn S, Sudbrak R, Albrecht MW, Lienhard M, Craig DW, Izatt T, Kurdoglu AA, Sherry ST, Ananiev V, Belaia Z, Beloslyudtsev D, Bouk N, Chen C, Church D, Cohen R, Cook C, Garner J, Hefferon T, Kimelman M, Liu C, Lopez J, Meric P, O’Sullivan C, Ostapchuk Y, Phan L, Ponomarov S, Schneider V, Shekhtman E, Sirotkin K, Slotta D, Xiao C, Zhang H, Haussler D, Abecasis GR, McVean GA, Alkan C, Ko A, Dooling D, Durbin RM, Balasubramaniam S, Keane TM, McCarthy S, Stalker J, Chakravarti A, Knoppers BM, Abecasis GR, Barnes KC, Beiswanger C, Burchard EG, Bustamante CD, Cai H, Cao H, Durbin RM, Gharani N, Gibbs RA, Gignoux CR, Gravel S, Henn B, Jones D, Jorde L, Kaye JS, Keinan A, Kent A, Kerasidou A, Li Y, Mathias R, McVean GA, Moreno-Estrada A, Ossorio PN, Parker M, Reich D, Rotimi CN, Royal CD, Sandoval K, Su Y, Sudbrak R, Tian Z, Timmermann B, Tishkoff S, Toji LH, Tyler-Smith C, Via M, Wang Y, Yang H, Yang L, Zhu J, Bodmer W, Bedoya G, Ruiz-Linares A, Ming CZ, Yang G, You CJ, Peltonen L, Garcia-Montero A, Orfao A, Dutil J, Martinez-Cruzado JC, Oleksyk TK, Brooks LD, Felsenfeld AL, McEwen JE, Clemm NC, Duncanson A, Dunn M, Green ED, Guyer MS, Peterson JL.

## Supplemental Methods

### Identifying STR loci in the human genome

#### Finding putative STR loci in the human genome

To identify putative STRs, we relied on the heuristic that these sequences should rarely occur in randomly generated sequences. To accomplish this aim, we determined the dinucleotide transition frequencies for each of the 22 autosomal chromosomes and 2 sex chromosomes. Then, for each chromosome, we used a second order Markov process to generate a random DNA sequence with the same length and transition frequencies. We repeated this process ten times for each chromosome. Next, Tandem Repeats Finder (TRF) (Benson 1999) was run on the random and real human chromosomes with a match weight of 2, a mismatch and indel penalty of 7, an 80% probability of matching and a 10% probability of an indel. In cases where TRF reported two overlapping STRs, we selected the locus with the highest score. For each chromosome, the repeats for periods 2-6bp were then analyzed separately to calculate the minimum TRF score where the number of repeats in the hg19 chromosomal sequence was at least 100 times greater than the mean number in the corresponding ten simulated chromosomes (Supplemental Figure 1a). These per-chromosome thresholds were combined into five genome-wide thresholds by taking the maximum threshold across all chromosomes (Supplemental Table 12). The identified repeats in hg19 were then filtered to only include those loci with a score above the threshold for its period. Finally, loci that originally overlapped other TRF results were removed, as these loci were generally very impure.

#### Incorporation of annotated STRs

To collate a set of annotated STR markers, we downloaded published PCR primers for the Marshfield markers from http://www.stanford.edu/group/rosenberglab/repeatsDownload.html (Pemberton et al. 2009.) We then utilized *in silico* PCR to determine the genomic coordinates of these primers. Locations for Y-STRs were obtained as described in our previous work (Gymrek et al. 2013) and locations for the CODIS markers were determined using PCR primers contained in the NIST database and the method outlined above. To integrate these markers into our reference set, we utilized BEDTools (Quinlan and Hall 2010) to remove any empirical loci that overlapped annotated loci. The remaining empirical loci combined with all annotated markers comprised our final STR reference.

#### Assessment of empirical score thresholds with a permissive call set

Having assembled a genome-wide STR reference, we sought to ensure that the chosen TRF score thresholds were appropriate. As dynamic expansions and contractions are hallmarks of STR loci, we examined the rates of polymorphism rates a function of TRF score in a permissive call set generated using data from the 1000 Genomes.

To this end, we created a permissive reference by running TRF using relaxed parameters. In particular, we ran the program using a much lower score cutoff of 14 (instead of a score greater than equal to the empirical TRF thresholds) to identify a candidate set of STRs with repeat unit sizes of 2 to 5bp. The other parameters used to run TRF were match = 2, indel = 5, mismatch = 5, match probability = 80, indel probability = 10, and minimum score = 14. We removed STRs that localized to areas that might preclude unique mapping, such as large repeats or transposable elements. Transposons and other repetitive elements were identified using RepeatMasker and the TRF results in or within 20 bases of these regions were removed. We further removed any STRs that were located next to or within 20 bases of another STR. Finally, we pruned the list of STRs on the basis of empirically derived tract length thresholds and purity thresholds, developed in a previous study that characterized the minimum requirements of a sequence to mutate as an STR (Fondon JW et al., 2012), namely minimum tract lengths of 13, 20, 23, and 27bp for 2-5mers, respectively.

We then loaded this permissive reference to RepeatSeq and generated a preliminary STR call set using data from Phase 1 of the 1000 Genomes project.

Encouragingly, we found that loci around the cutoffs identified by running TRF on the random chromosomes were close to fixation. The mean heterozygosity rapidly increased shortly after the threshold (Supplemental Figure 1b). This phenomenon matches the hallmark of mature STRs that dynamically expand and contract.

We further quantified the number of polymorphic loci (heterozygosity > 2%) omitted and included by these score thresholds (Supplemental Table 1). These analyses revealed that less than 1% of loci omitted by the thresholds were polymorphic, while roughly 40% of included loci were polymorphic. In addition, our thresholds only omitted 1% of all polymorphic loci. Collectively, these analyses strongly suggest that our score thresholds are well calibrated and have a low false negative rate.

### Call set integration

RepeatSeq (Highnam et al. 2012) and lobSTR (Gymrek et al. 2012) are currently the two primary tools utilized to genotype STRs in high-throughput sequencing data. As a result, to create the most accurate call set, we sought to assess their individual performance and potentially integrate their calls if it improved accuracy. lobSTR calls were generated as previously described (**Methods**) while RepeatSeq (available http://github.com/adaptivegenome/repeatseq) was run using default parameters on the read alignments produced by the 1000 Genomes project.

To assess call set performance, we utilized the Marshfield capillary electrophoresis (CE) genotypes generated for a subset of the 1000 Genomes samples (Rosenberg et al. 2005). The loci in this panel are highly polymorphic and provide a challenging test set to compare between lobSTR and RepeatSeq. We regressed the dosage produced by the method of interest (the sum of the predicted base pair differences from the reference allele) versus the dosage obtained from CE (Supplemental Figure 2A-B) Comparing the calls produced by the individual methods indicated that lobSTR outperformed RepeatSeq as the R^2^ values were 0.71 and 0.4, respectively. While RepeatSeq produced more calls, they were in general strongly biased towards the reference allele genotype.

To integrate calls from RepeatSeq and lobSTR, we first had to combine the genotype likelihoods of lobSTR and RepeatSeq. While RepeatSeq reports *P*(*genotype*|*data*), lobSTR reports *P*(*genotype*|*data*) Therefore, we generated comparable posteriors for the lobSTR calls by using the population-wide occurrences of each STR allele in the lobSTR VCF “GT” field as a prior.

In total, we explored three different simplistic integration strategies. The first strategy selected the genotype with the highest posterior likelihood across both methods. This method assumes that the lobSTR and RepeatSeq posteriors are well calibrated and therefore defers to the method with the most confidence. We explored two variants of this strategy by considering a) only calls for which both methods produced genotypes and b) calls for which either method produced genotypes. The resulting R^2^ values of 0.52 and 0.53 (Supplemental Figure 2C-D) indicated that the integrated calls outperformed those of RepeatSeq but were still inferior to those of lobSTR alone. The next integration strategy selected the genotype with the highest mean posterior. When we applied this strategy in combination with the two aforementioned variants, the R^2^ values were nearly identical to those obtained by selecting the maximum posterior (Supplemental Figure 2E-F). Finally, we employed a simple strategy in which we only considered concordant calls. In addition to greatly limiting the number of calls, this strategy was surprisingly worse than the previous two integration strategies as it resulted in an of R^2^ 0.46 (Supplemental Figure 2G).

In summary, despite various attempts to integrate RepeatSeq and lobSTR calls, our efforts were ultimately unsuccessful. Though the integrated calls were superior to those of RepeatSeq alone, they were ultimately inferior to lobSTR's calls. The results of these efforts suggested that RepeatSeq calls had systematic biases and that these biases, when integrated with lobSTR calls, persisted. We therefore chose to proceed using only the lobSTR calls.

**Supplemental Figure 1:**
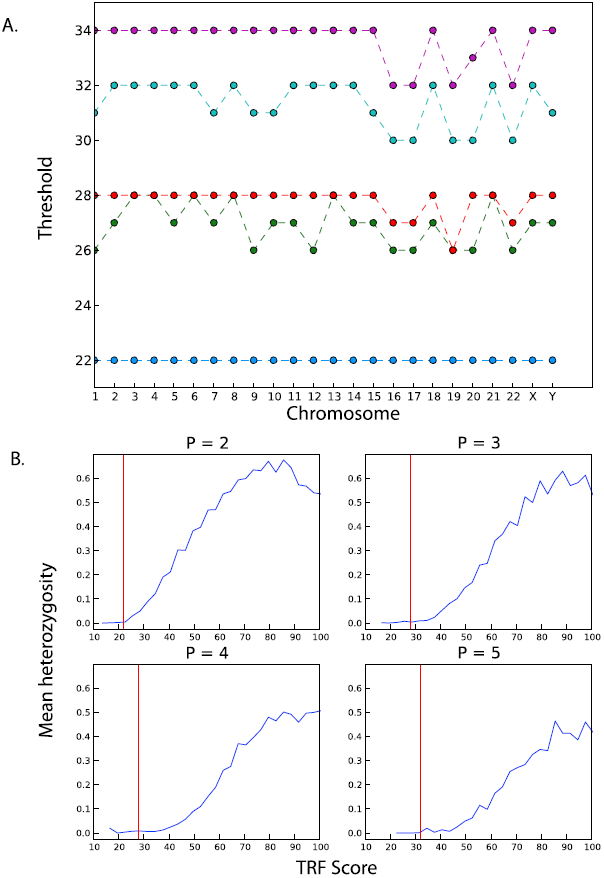
Evidence-based criteria for STR loci. (A) The TRF score threshold for each chromosome and motif length. These thresholds were calculated based on an FDR of 1% using the second-order Markov chain simulations (blue: 2mer, green: 3mer, red: 4mer, cyan: 5mer, purple: 6mer). **(B) Validating the thresholds with a preliminary call set.** The plots show the average heterozygosity levels (y-axis) for STRs in the permissive catalog as a function of their TRF scores (x-axis). P denotes the motif length in bp. The red line shows the thresholds that were selected for the final definition based on the Markov chain simulations. The putative STRs around the thresholds are close to fixation and STRs with TRF score above the threshold show a rapid increase in their heterozygosity. This indicates that the thresholds are well calibrated and include most of the STRs that are subject to contractions and expansions.

**Supplemental Figure 2:**
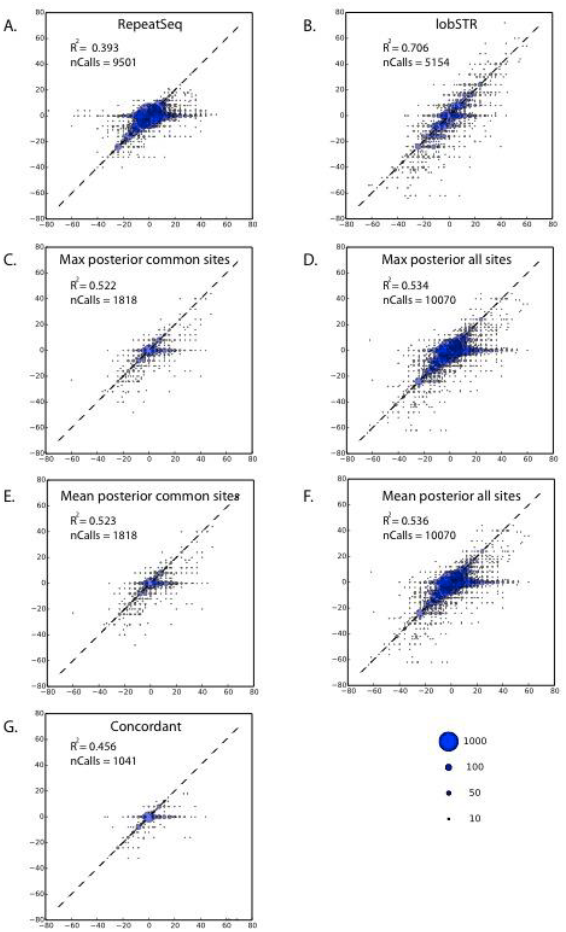
Integration efforts for RepeatSeq and lobSTR. Each bubble plot shows the regression of the Marshfield capillary dosages (x-axis) with a different method to obtain STR calls from the 1000 Genomes (y-axis). The R^2^ and number of calls (nCalls) are reported **(A) RepeatSeq alone (B) lobSTR alone (C) RepeatSeq + lobSTR integration based on maximum posterior for sites that appeared in both datasets (D) RepeatSeq + lobSTR integration based on maximum posterior for sites that appeared in at least one dataset (E) RepeatSeq + lobSTR based on mean posterior for sites that appeared in both datasets (F) RepeatSeq + lobSTR based on mean posterior for sites that appeared in at least one dataset(G) RepeatSeq + lobSTR integration by reporting only genotypes concordant between the two methods.** The best R^2^ was obtained by lobSTR alone.

**Supplemental Figure 3:**
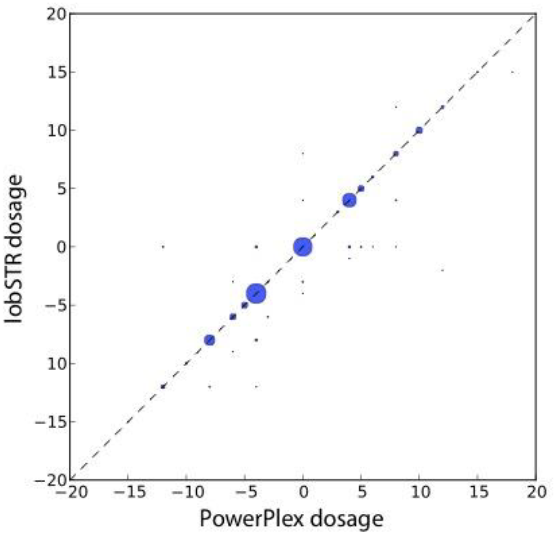
lobSTR dosage concordance with capillary electrophoresis for hemizygous Y-STRs. The dosage is reported as the base pair difference from the NCBI reference. The area of each bubble is proportional to the number of calls of the dosage combination and the broken line indicates the diagonal.

**Supplemental Figure 4:**
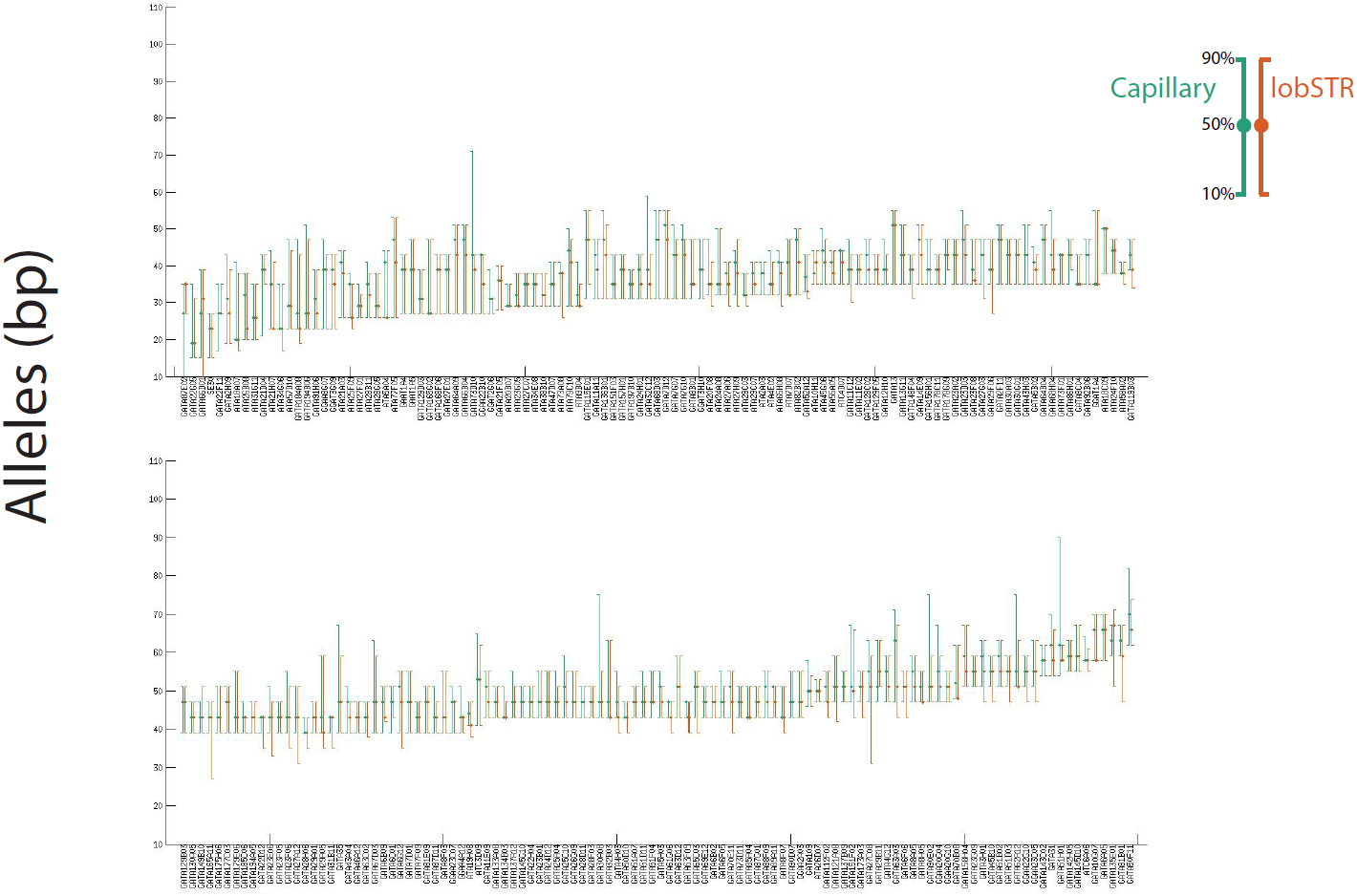
Allelic spectra comparisons of capillary results (green) versus lobSTR (orange) for the Marshfield panel. Each rod represents the interdecile range of the STR alleles in the European population (in bp) and the median allele.

**Supplemental Figure 5:**
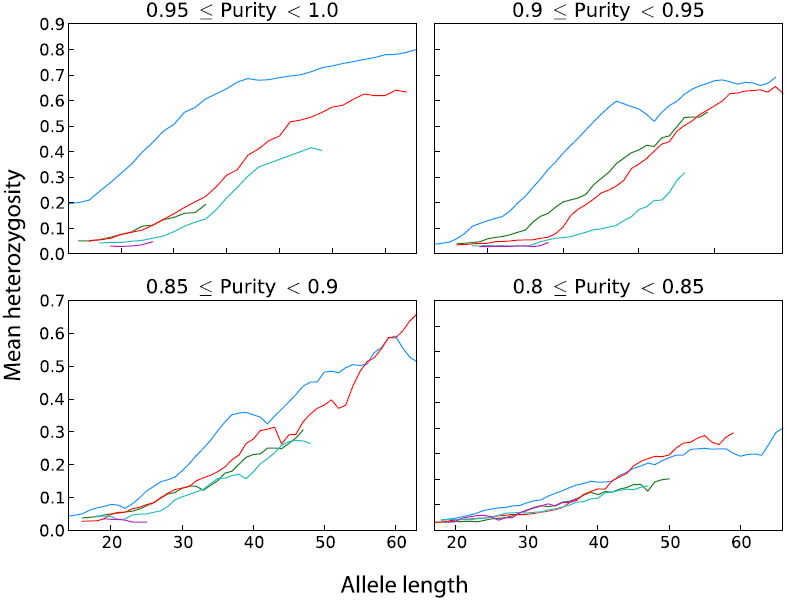
STR variability as a function of wild-type allele length (bp) for impure STR loci. Analysis is stratified based on motif length (blue: 2mer, green: 3mer, red: 4mer, cyan: 5mer, purple: 6mer) and the purity of the STR (see methods) and is restricted to STRs whose wild-type allele matches the reference.

**Supplemental Figure 6:**
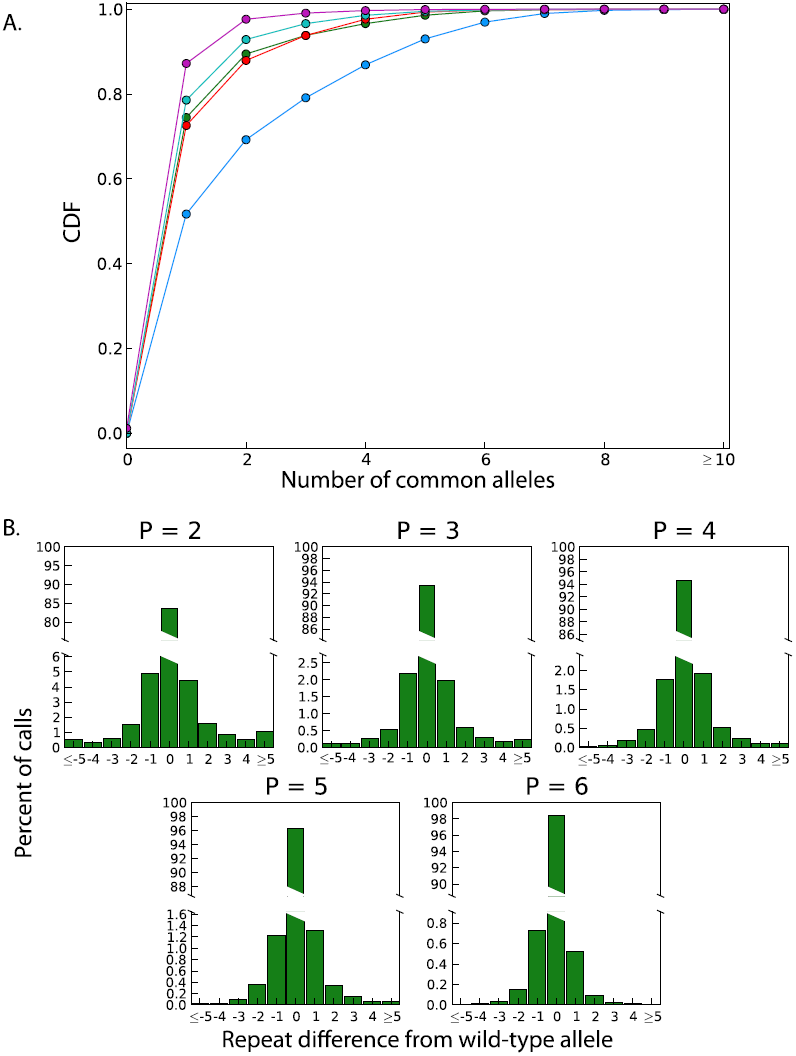
Patterns of STR variation. **(A) The cumulative distribution function of the number of alleles with MAF > 5% stratified by motif length** (blue: 2mer, green: 3mer, red: 4mer, cyan: 5mer, purple: 6mer) **(B) The averaged allelic spectra of STRs.** P denotes the motif length in bp. The 0 allele is the most common allele (wild-type allele).

**Supplemental Figure 7:**
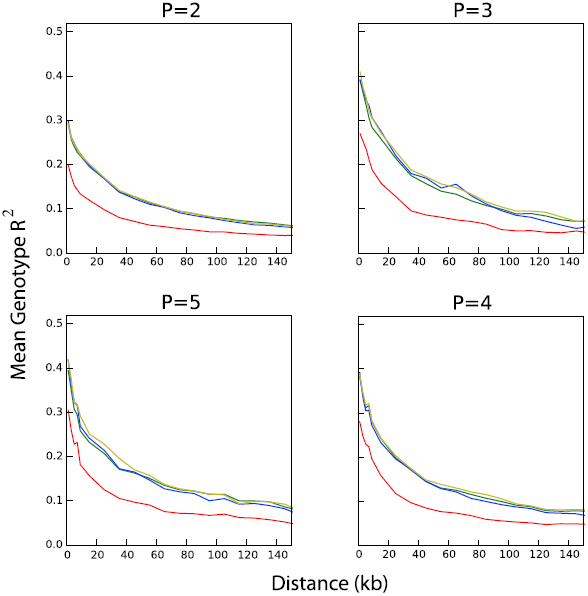
STR-SNP linkage disequilibrium on chromosome X stratified by motif length. P denotes the motif length in bp. Africans (red), Admixed Americans (green), Europeans (yellow) and East Asians (blue). Longer repeat motifs show an increase in the level of STR-SNP LD.

**Supplemental Table 1.**
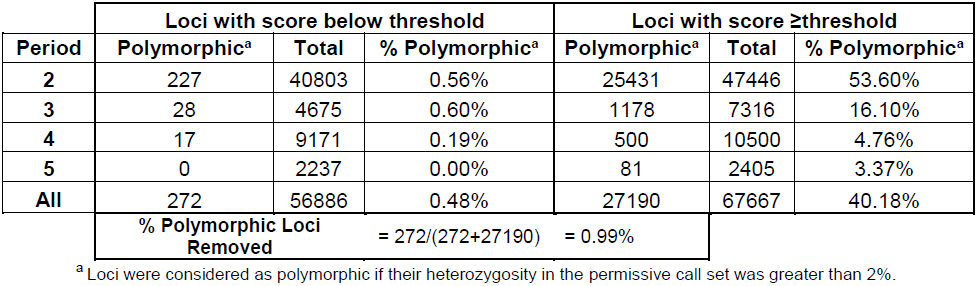
Polymorphism levels of loci omitted and included by TRF score cutoffs.

**Supplemental Table 2.** The distribution of STRs in the lobSTR reference.

| **Period** | **Total STRs** |
| --- | --- |
| **2** | 277822 |
| **3** | 77327 |
| **4** | 220859 |
| **5** | 72637 |
| **6** | 40867 |
| **Total** | 689512 |

**Supplemental Table 3.** Population breakdown of genotyped samples.

| **Population Abbreviation** | **Population Name** | **Continent** | **Super group** | **Number of Individuals** |
| --- | --- | --- | --- | --- |
| ASW | Americans of African Ancestry in SW USA | North America | African | 50 |
| CEU | Utah Residents with Northern and Western European Ancestry | Europe | European | 74 |
| CHB | Han Chinese in Beijing, China | Asia | East Asian | 82 |
| CHS | Southern Han Chinese | Asia | East Asian | 98 |
| CLM | Columbians from Medellin, Colombia | South America | Admixed American | 59 |
| FIN | Finnish in Finland | Europe | European | 87 |
| GBR | British in England and Scotland | Europe | European | 79 |
| IBS | Iberian population in Spain | Europe | European | 16 |
| JPT | Japanese in Tokyo, Japan | Asia | East Asian | 82 |
| LWK | Luhya in Webuye, Kenya | Africa | African | 82 |
| MXL | Mexican Ancestry from Los Angeles | North America | Admixed American | 58 |
| PUR | Puerto Ricans from Puerto Rico | North America | Admixed American | 70 |
| TSI | Toscani in Italia | Europe | European | 89 |
| YRI | Yoruba in Ibadan, Nigeria | Africa | European | 83 |
| Total |  |  |  | 1009 |

**Supplemental Table 4.** Marshfield concordance statistics.

|  | **lobSTR**<br>**A/A Call** | **lobSTR**<br>**C/C Call** | **lobSTR**<br>**A/B Call** | **lobSTR**<br>**A/C Call** | **lobSTR C/D**<br>**Call** |
| --- | --- | --- | --- | --- | --- |
| **Homozygous**<br>**Marshfield**<br>**Sites (A/A)** | 1465<br>(89.5%) | 119<br>(7.3%) | N/A | 52<br>(3.2%) | 1 (< .1%) |
| **Heterozygous**<br>**Marshfield**<br>**Sites (A/B)** | 2681<br>(76.0%) | 290<br>(8.2%) | 452<br>(12.8%) | 95<br>(2.7%) | 9 (.3%) |

Marshfield genotypes are always A/A or A/B.
lobSTR calls with C and/or D alleles indicate alleles that were not observed in the Marshfield genotype.

**Supplemental Table 5.** Y-STR PowerPlex concordance statistics.

| **Locus** | **Correct Calls** | **Total Calls** | **% Correct** |
| --- | --- | --- | --- |
| DYS481 | 14 | 19 | 73.68 |
| DYS458 | 10 | 12 | 83.33 |
| DYS533 | 60 | 66 | 90.91 |
| DYS389I | 53 | 58 | 91.38 |
| DYS392 | 36 | 39 | 92.31 |
| DYS391 | 98 | 105 | 93.33 |
| DYS456 | 47 | 50 | 94.00 |
| DYS438 | 51 | 53 | 96.23 |
| DYS393 | 57 | 59 | 96.61 |
| DYS439 | 83 | 85 | 97.65 |
| Y-GATA-H4 | 49 | 50 | 98.00 |
| DYS549 | 66 | 67 | 98.51 |
| DYS19 | 15 | 15 | 100.00 |
| DYS437 | 14 | 14 | 100.00 |
| DYS570 | 15 | 15 | 100.00 |
| DYS576 | 26 | 26 | 100.00 |
| DYS643 | 51 | 51 | 100.00 |
| **Total** | **745** | **784** | **95.03** |

**Supplemental Table 6.** Comparing heterozygosities between populations.

|  |  | Population B |  |  |  |  |  |  |  |  |  |
| --- | --- | --- | --- | --- | --- | --- | --- | --- | --- | --- | --- |
|  |  | CEU | FIN | GBR | TSI | ASW | LWK | YRI | CHB | CHS | JPT |
| Population A | CEU |  | 0.53 | 0.46 | 0.51 | 0.36 | 0.39 | 0.4 | 0.55 | 0.55 | 0.53 |
|  | FIN | 0.47 |  | 0.43 | 0.48 | 0.35 | 0.38 | 0.38 | 0.53 | 0.53 | 0.5 |
|  | GBR | 0.54 | 0.57 |  | 0.55 | 0.38 | 0.41 | 0.41 | 0.57 | 0.57 | 0.55 |
|  | TSI | 0.49 | 0.52 | 0.45 |  | 0.36 | 0.39 | 0.39 | 0.54 | 0.54 | 0.52 |
|  | ASW | 0.64 | 0.65 | 0.62 | 0.64 |  | 0.53 | 0.54 | 0.68 | 0.67 | 0.66 |
|  | LWK | 0.61 | 0.62 | 0.59 | 0.61 | 0.47 |  | 0.51 | 0.65 | 0.64 | 0.63 |
|  | YRI | 0.6 | 0.62 | 0.59 | 0.61 | 0.46 | 0.49 |  | 0.64 | 0.64 | 0.62 |
|  | CHB | 0.45 | 0.47 | 0.43 | 0.46 | 0.32 | 0.35 | 0.36 |  | 0.5 | 0.46 |
|  | CHS | 0.45 | 0.47 | 0.43 | 0.46 | 0.33 | 0.36 | 0.36 | 0.5 |  | 0.46 |
|  | JPT | 0.47 | 0.5 | 0.45 | 0.48 | 0.34 | 0.37 | 0.38 | 0.54 | 0.54 |  |

Only the 10% most variable STRs were used. Each cell represents the fraction of loci for which population “A” had higher heterozygosity than population “B”.

**Supplemental Table 7.**
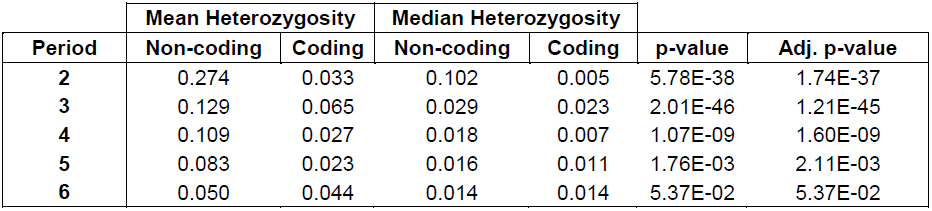
Comparison of heterozygosity in noncoding and coding STR.

| **Period** | **Mean Heterozygosity** |  | **Median Heterozygosity** |  | **p-value** | **Adj. p-value** |
| --- | --- | --- | --- | --- | --- | --- |
|  | **Non-coding** | **Coding** | **Non-coding** | **Coding** |  |  |
| **2** | 0.274 | 0.033 | 0.102 | 0.005 | 5.78E-38 | 1.74E-37 |
| **3** | 0.129 | 0.065 | 0.029 | 0.023 | 2.01E-46 | 1.21E-45 |
| **4** | 0.109 | 0.027 | 0.018 | 0.007 | 1.07E-09 | 1.60E-09 |
| **5** | 0.083 | 0.023 | 0.016 | 0.011 | 1.76E-03 | 2.11E-03 |
| **6** | 0.050 | 0.044 | 0.014 | 0.014 | 5.37E-02 | 5.37E-02 |

**Supplemental Table 8.**
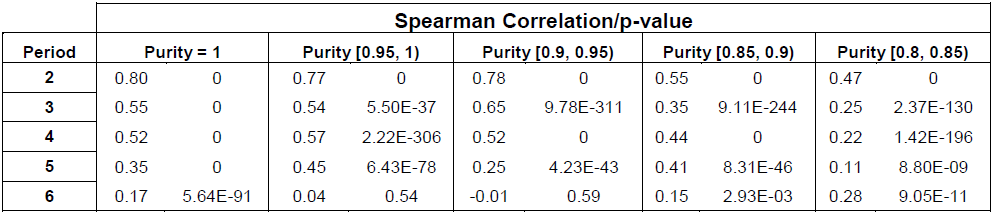
Correlation between allele length and heterozygosity for various levels of STR purity.

| Period | Spearman Correlation/p-value |  |  |  |  |  |  |  |  |  |
| --- | --- | --- | --- | --- | --- | --- | --- | --- | --- | --- |
|  | Purity = 1 |  | Purity [0.95, 1) |  | Purity [0.9, 0.95) |  | Purity [0.85, 0.9) |  | Purity [0.8, 0.85) |  |
| **2** | 0.80 | 0 | 0.77 | 0 | 0.78 | 0 | 0.55 | 0 | 0.47 | 0 |
| **3** | 0.55 | 0 | 0.54 | 5.50E-37 | 0.65 | 9.78E-311 | 0.35 | 9.11E-244 | 0.25 | 2.37E-130 |
| **4** | 0.52 | 0 | 0.57 | 2.22E-306 | 0.52 | 0 | 0.44 | 0 | 0.22 | 1.42E-196 |
| **5** | 0.35 | 0 | 0.45 | 6.43E-78 | 0.25 | 4.23E-43 | 0.41 | 8.31E-46 | 0.11 | 8.80E-09 |
| **6** | 0.17 | 5.64E-91 | 0.04 | 0.54 | -0.01 | 0.59 | 0.15 | 2.93E-03 | 0.28 | 9.05E-11 |

**Supplemental Table 9.** Allele frequency distributions for STR type subsets.

| Period | Number of Loci | Length Percentile (%) | Wild-type Allele Length (bp)<sup>a</sup> | Wild-type Allele Frequency<sup>a,b</sup> | MAF<sup>a,b,c</sup> |
| --- | --- | --- | --- | --- | --- |
| **2** | 253,809 | 0-25 | 12 | 97.95 | 1.17 |
|  |  | 25-50 | 16 | 96.44 | 2.17 |
|  |  | 50-75 | 25 | 88.64 | 7.02 |
|  |  | 75-100 | 38 | 53.52 | 16.03 |
| **3** | 71,582 | 0-25 | 15 | 98.85 | 0.51 |
|  |  | 25-50 | 18 | 98.67 | 0.67 |
|  |  | 50-75 | 22 | 98.28 | 0.94 |
|  |  | 75-100 | 32 | 95.64 | 2.68 |
| **4** | 195,264 | 0-25 | 14 | 99.34 | 0.27 |
|  |  | 25-50 | 18 | 99.24 | 0.31 |
|  |  | 50-75 | 23 | 98.96 | 0.49 |
|  |  | 75-100 | 35 | 95.72 | 2.73 |
| **5** | 64,320 | 0-25 | 17 | 99.33 | 0.27 |
|  |  | 25-50 | 20 | 99.25 | 0.30 |
|  |  | 50-75 | 24 | 99.16 | 0.36 |
|  |  | 75-100 | 34 | 98.60 | 0.75 |
| **6** | 30,063 | 0-25 | 17 | 99.30 | 0.28 |
|  |  | 25-50 | 21 | 99.31 | 0.28 |
|  |  | 50-75 | 23 | 99.30 | 0.29 |
|  |  | 75-100 | 32 | 99.24 | 0.36 |

<sup>a</sup> The reported number is the median of all values within the length percentile bin.
<sup>b</sup> Frequencies are given as percentages.
<sup>c</sup> The frequency of the second most common STR allele.

**Supplemental Table 10.**
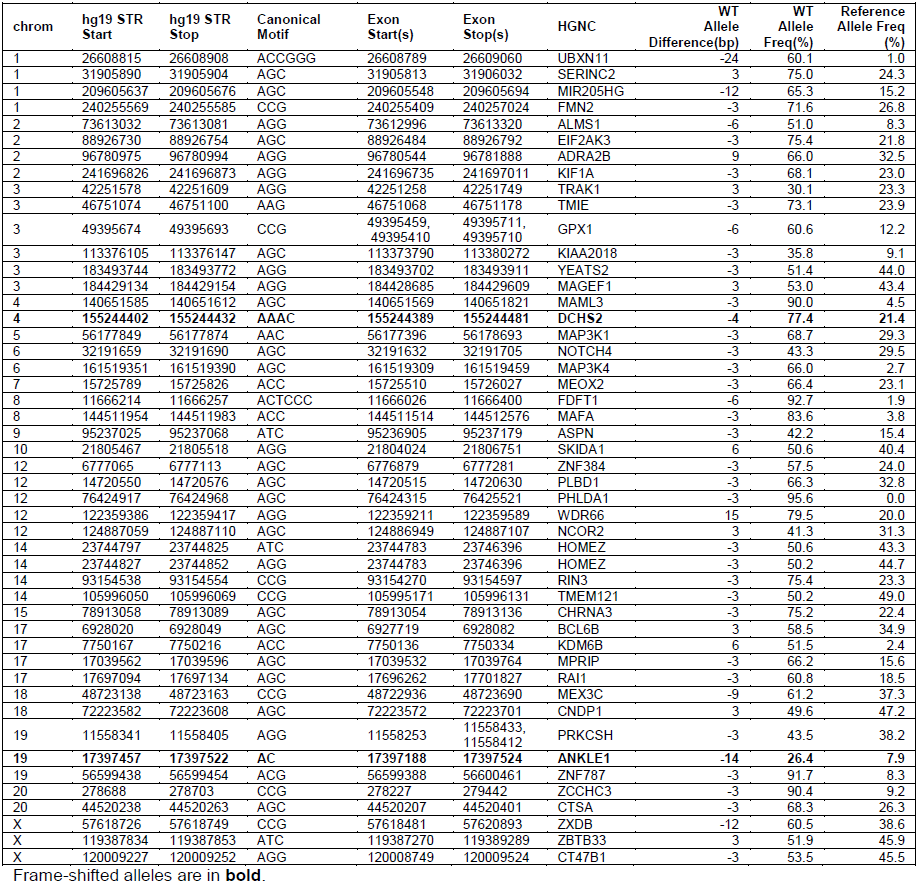
Coding STRs with non-reference wild-type alleles.

**Supplemental Table 11.** Common LoF alleles.

| chr | STR start | STR stop | bp from hg19 | Motif | # Samples | Exon Start | Exon Stop | Gene |
| --- | --- | --- | --- | --- | --- | --- | --- | --- |
| 4 | 155244402 | 155244432 | -4 | AAAC | 395 | 155244389 | 155244481 | DCHS2 |
| 9 | 35561913 | 35561938 | -8 | ACCC | 22 | 35561864 | 35562092 | FAM166B |
| 10 | 81841429 | 81841443 | -4 | AAAG | 25 | 81841395 | 81841490 | TMEM254 |
| 10 | 81841429 | 81841443 | 1 | AAAG | 111 | 81841395 | 81841490 | TMEM254 |
| 10 | 81841429 | 81841443 | 2 | AAAG | 31 | 81841395 | 81841490 | TMEM254 |
| 19 | 55526092 | 55526121 | 4 | ACAG | 82 | 55525449 | 55526533 | GP6 |
| 20 | 48467310 | 48467334 | -1 | AC | 94 | 48467298 | 48467381 | SLC9A8 |

**Supplemental Table 12.** TRF score cutoffs.

| **Period** | **Score Cutoff** |
| --- | --- |
| 2 | 22 |
| 3 | 28 |
| 4 | 28 |
| 5 | 32 |
| 6 | 34 |

